# Interaction with the membrane-anchored protein CHIC2 constrains the ubiquitin ligase activity of CHIP

**DOI:** 10.1101/2023.07.17.549407

**Authors:** M.D. Callahan, M. Hodul, E.C. Carroll, M. Ravalin, C.M. Nadel, A.R.I. de Silva, A.R. Cupo, L.E. McDermott, J.C. Nix, R.C. Page, A.W. Kao, J.E. Gestwicki

## Abstract

Maintenance of cellular health requires the proper regulation of E3 ubiquitin ligases. The E3 ligase CHIP is canonically regulated by its interactions with the molecular chaperones Hsp70 and Hsp90, which focus CHIP’s ubiquitination activity on misfolded proteins. Here, we report a chaperone-independent interaction of CHIP with the membrane-anchored protein CHIC2, which strongly attenuates CHIP’s ligase activity. We show that CHIC2 outcompetes abundant, cytosolic chaperones through its exquisite CHIP selectivity, rather than through enhanced affinity. In proteomic experiments, we find that CHIC2 knockout phenocopies CHIP knockout in certain cell types, implying that chaperone-independent interactions can sometimes predominate CHIP’s functions. Furthermore, loss of the CHIP-CHIC2 interaction induces neurodegeneration and shortens lifespan in *C. elegans*, demonstrating that formation of this chaperone-independent complex is important in animals. We propose that CHIC2 attenuates CHIP activity at the membrane, offering a novel mechanism by which this ubiquitin ligase can be regulated.

## Introduction

Ubiquitination regulates a wide range of protein functions, including localization, activity and turnover^1^. This post-translational modification (PTM) is achieved through the coordinated action of E1, E2 and E3 ubiquitin ligases, which recognize specific substrates and transfer mono- or poly-ubiquitin chains^2^. There is a long-standing interest in deciphering the regulatory networks responsible for controlling ubiquitination. For example, many ubiquitinated proteins are known to be further processed by the successive action of multiple E3 ligases and/or de-ubiquitinating enzymes (DUBs), which refine the length or connectivity of the ubiquitin chains^3^. The relative activity and substrate selectivity of the E3 ligases themselves are also regulated, through processes such as PTMs^4^ and changes in oligomeric state^5^. Finally, modulation of E3 ligase activity through specific protein-protein interactions is also possible, as illustrated by proteins such as Emi1, which regulates the cell cycle by acting as a pseudosubstrate inhibitor of the anaphase promoting complex/cyclosome (APC/C)^6^. Together, these observations illustrate the wide variety of mechanisms by which protein ubiquitination can be regulated.

The carboxy-terminus of Hsp70 interacting protein (CHIP) is an E3 ubiquitin ligase that is canonically associated with polyubiquitination and turnover of damaged or terminally misfolded proteins^7^. CHIP is highly expressed in the brain, and its activity is particularly important for the degradation of proteins important in neurodegenerative diseases^8^. CHIP is composed of a coiled-coil domain that mediates dimerization, a U-box domain that recruits E2 conjugating enzymes, and a tetratricopeptide repeat (TPR) domain. The TPR domain of CHIP is known to bind a conserved EEVD motif that is located at the extreme C-termini of the cytosolic chaperones heat shock protein 70 (Hsp70) and heat shock protein 90 (Hsp90)^9^. Crystal structures of EEVD-containing peptides bound to CHIP’s TPR domain have shown that key molecular contacts include coordination of the aspartate and terminal carboxylic acids by cationic residues in CHIP’s TPR domain, termed the “two-carboxylate clamp”^10^. This protein-protein interaction (PPI) is believed to recruit CHIP to misfolded proteins, with the chaperones acting as adapters to localize CHIP’s activity. However, recent work has identified other proteins that contain EEVD-like motifs^11,12^, suggesting that chaperones are not the only partners of CHIP (**Fig 1a)**. Moreover, CHIP also recognizes some substrates, such as interferon regulatory factor 1 (IRF1) and microtubule-associated protein tau (MAPT), independent of a chaperone or EEVD-like motif^13,14^. Together, these observations suggest that CHIP has a wider set of potential partners than previously anticipated. However, Hsp70s and Hsp90s are highly abundant proteins, so the relative contributions of chaperone-dependent and chaperone-independent mechanisms are not yet clear.

**Figure 1:**
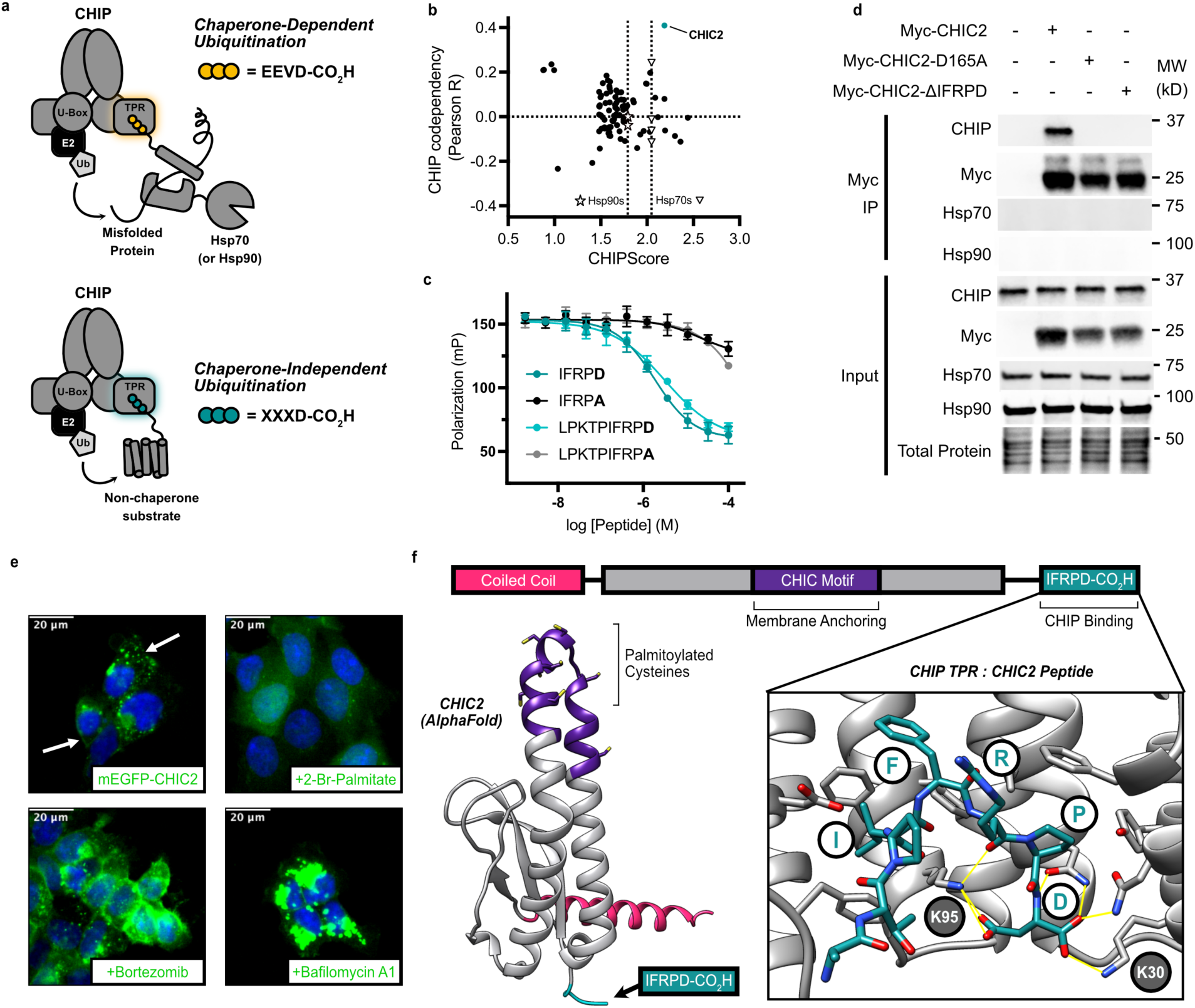
Direct C-terminal binding mediates CHIP’s interaction with the membrane-anchored protein CHIC2. **a)** Schematic comparing the canonical, chaperone-dependent mechanism of CHIP to the chaperone-independent mechanism. Both processes are rooted in recognition of C-terminal aspartic acids within EEVD-like motifs. **b)** A combined ranking of putative CHIP substrates using CHIPscore and genetic codependency from the Cancer DepMap reveals a functional interaction between CHIC2 and CHIP. For reference, CHIP’s canonical partners, Hsp70 (open triangles) and Hsp90 (open stars), are highlighted. **c)** Competitive displacement of a fluorescence polarization tracer from CHIP’s TPR domain by 5 or 10 amino acid peptides derived from CHIC2’s C-terminus. Mutation of the C-terminal aspartate abolishes binding. Error bars represent S.D. of 4 replicate datapoints from 1 of 3 representative experiments. **d)** Co-immunoprecipitation of CHIC2 and CHIP from HEK293T cells. Data is representative of 3 independent biological experiments. **e)** Live-cell fluorescence microscopy of dox-inducible Flp-In T-REX HEK-293 cells expressing mEGFP-CHIC2. CHIC2 localization to the plasma membrane and vesicles is shown (arrows). Inhibitors: 2-bromo-palmitate (palmitoylation inhibitor); bortezomib (proteasome inhibitor); bafilomycin A1 (lysosome inhibitor). **f)** CHIC2 domain architecture and AlphaFold2 predicted structure, highlighting the membrane-binding CHIC domain and the EEVD-like, C-terminal motif. Co-crystal structure of the CHIC2 peptide bound to CHIP’s TPR domain (inset; 1.63 Å resolution) reveals the expected interactions between the terminal aspartate and the two-carboxylate clamp residues K30 and K95 (also see Extended Data 1).

To better understand what other proteins might bind to CHIP, we recently screened large-scale peptide libraries that resemble the EEVD motif^11^. This screen revealed CHIP’s preference for each amino acid within the five C-terminal positions of a substrate, resulting in an algorithm, termed CHIPScore, that predicts the affinity of short peptide motifs for CHIP’s TPR domain. To test this algorithm, we used it to design an optimized ligand, termed CHIPOpt (Ac-LWWPD), that binds 50-fold tighter than the Hsp70 and Hsp90 sequences. Here, we reasoned that CHIPScore might also be used to search for potential non-canonical CHIP partners in the human genome. Specifically, we hypothesized that other proteins might contain previously unrecognized, EEVD-like motifs that would bind CHIP and potentially serve as chaperone-independent regulators of its E3 ligase functions. Indeed, by combining CHIPScore with functional genomics data in the Cancer Dependency Map (DepMap), we report here that the relatively uncharacterized, membrane-anchored protein CHIC2 is a strong and biologically important binding partner of CHIP. We show that CHIC2 contains an EEVD-like motif that binds to CHIP *in vitro* and in cells, and that this interaction is chaperone-independent. We find that CHIC2 binding dramatically reduces CHIP ubiquitin ligase activity *in vitro*, suggesting that it locally restricts CHIP function at the cell surface and vesicles where CHIC2 is typically localized. Using a chemoproteomic assay, we show that CHIC2 effectively out-competes the abundant chaperones Hsp70 and Hsp90, seemingly because of its exquisite selectivity for CHIP over other TPR domain containing proteins. Moreover, using quantitative proteomics we show that loss of CHIC2 phenocopies CHIP knockout in some cells, but not others, suggesting that the CHIC2-CHIP axis may predominate over the canonical, chaperone-mediated mechanism in certain contexts. Finally, we show that the interaction between CHIC2 and CHIP is evolutionarily conserved, and that disrupting CHIC2’s EEVD-like motif induces premature aging and neurodegeneration in *C. elegans.* We propose that CHIC2 is an important, chaperone-independent regulator of CHIP function, and that its activity sometimes predominates over CHIP’s canonical, chaperone-mediated mechanisms.

## Results

### Direct C-terminal binding mediates CHIP’s interaction with the membrane-anchored protein CHIC2

We previously developed a biophysics-based scoring function, termed CHIPScore, that predicts the affinity of CHIP’s TPR domain for putative EEVD-like sequences at the extreme C-terminus of open reading frames (ORFs)^11^. To prioritize biologically important hits for further study, we searched the DepMap database^15^ for genes that shared a genetic co-dependency with CHIP and then ranked the C-terminal sequences of those proteins using CHIPScore. We hypothesized that this approach would focus on the most biologically important, direct interactors of CHIP. This search process revealed a striking relationship with the relatively uncharacterized protein, CHIC2, which was a significant outlier in both its genetic interaction and predicted binding affinity for CHIP (**Fig. 1b**).

To test whether CHIC2’s C-terminus might indeed bind to CHIP’s TPR domain *in vitro*, we tested peptides corresponding to either the last five (Ac-IFRPD) or ten (Ac-LPKTPIFRPD) amino acids of CHIC2 and monitored their binding to CHIP’s TPR domain using a fluorescence polarization (FP) assay. Consistent with a canonical carboxylate clamp binding mode, both of the peptides displaced a fluorescent tracer from CHIP’s TPR domain, while mutation of their C-terminal aspartic acids abolished binding (**Fig. 1c**). Next, to determine if the CHIC2-CHIP interaction also occurs in cells, we performed co-immunoprecipitation experiments. We expressed a myc-tagged CHIC2 protein in HEK293T cells and found that it bound endogenous CHIP. Importantly, mutation of the critical, C-terminal aspartic acid in CHIC2 (CHIC2^D165A^) was also sufficient to completely abolish binding (**Fig 1d**), re-enforcing the importance of CHIC2’s C-terminus in recruiting CHIP.

We also noticed that the immunoprecipitated complexes were depleted of Hsp70 and Hsp90, implying that CHIC2 displaces chaperone from both binding sites in an immunoprecipitated CHIP dimer. To understand how CHIC2’s C-terminus might compete with the canonical EEVD motifs of Hsp70 and Hsp90, we solved the crystal structure of the CHIC2 peptide (EFLPKTPIFRPD) bound to CHIP’s TPR domain (residues 21-154). This 1.63 Å co-structure confirmed interactions between the terminal aspartic acid and the two-carboxylate clamp residues in CHIP: K30 and K95 (**Fig. 1f, inset**).The overall binding pose of the CHIC2 peptide is highly similar to known CHIP ligands, such as the Hsp70 EEVD motif^16^ or CHIPOpt. For example, the phenylalanine and isoleucine in the CHIC2 peptide contact the “hydrophobic shelf” that is formed by F99, mimicking interactions made by the tryptophan and leucine in CHIPOpt. A feature unique to CHIC2 is an internal hydrogen bond between the arginine sidechain and peptide backbone that likely helps to enforce the bound conformation (**Extended Data 1**). Together, these findings confirm that CHIC2 has a validated EEVD-like motif that binds to CHIP’s TPR domain *in vitro* and in cells.

CHIC2 is named for its cysteine-rich hydrophobic (CHIC) domain, which is constitutively palmitoylated and known to localize CHIC2 to the plasma membrane and vesicles^17^. To validate this finding, we generated HEK293 Flp-In cells expressing mEGFP-CHIC2 under a doxycycline-inducible promoter. This platform allowed us to titrate CHIC2 expression to a minimal level (**Extended Data 2b**), limiting the potential for mislocalization due to overexpression. We then performed live-cell confocal imaging experiments, which revealed that CHIC2 is indeed localized to plasma membrane and vesicular compartments (**Fig 1e**). This localization disappeared upon treatment with the broad-spectrum palmitoylation inhibitor, 2-bromopalmitate (25 µM) (**Fig. 1e**), confirming the importance of this PTM in anchoring CHIC2 to the membrane. Deletion of CHIC2’s C-terminus (CHIC2-ΔIFRPD) did not substantially alter its subcellular localization but it did increase total levels of the protein (**Extended Data 2a**), suggesting that the interaction with CHIP is not required for localization but that it may be partially responsible for regulating CHIC2’s stability. Indeed, treatment with either bortezomib or bafilomycin increased CHIC2 levels (**Fig 1e**). However, we observed a more significant accumulation of total CHIC2 upon bafilomycin treatment (**Extended Data 2c**), suggesting that the protein is predominantly cleared through lysosomal degradation.

Given that the co-IP results showed that CHIC2 binds CHIP in cells, we hypothesized that the C-terminus of CHIC2 might be positioned away from the membrane, enabling it to recruit CHIP. An AlphaFold2 predicted structure of CHIC2 supports this idea, with the membrane-anchoring cysteines of the CHIC motif being positioned at the end of a helix-turn-helix, leaving the C-terminus exposed (**Fig. 1f**). To test whether CHIP can be directly recruited to the membrane by CHIC2, we attempted to observe their colocalization in the GFP-CHIC2 Flp-In cells. However, we did not observe substantial CHIP re-localization upon CHIC2 overexpression (**Extended Data 2d**). In contrast, CHIP over-expression did reduce CHIC2 levels and partially disrupted its vesicular localization, effects that required an intact CHIC2 C-terminus (**Extended Data 2d**). Interestingly, this effect only occurred in cells expressing CHIP at the highest levels (**Extended Data 2d).** Because both CHIC2 and cytosolic chaperones must compete for the same site on CHIP’s TPR domain, we speculate that chaperone-free CHIP levels may need to reach a certain threshold before CHIC2 can be substantially bound.

### The CHIC2 C-terminus outcompetes chaperone binding to CHIP using selectivity rather than affinity

What governs the competition between CHIC2 and chaperones for limited pools of CHIP? To better understand this process, we first compared the relative affinity of various C-terminal peptides using saturation FP assays. Designed peptides, such as CHIPOpt (LWWPD) and IWWPD^18^ bind CHIP’s TPR domain with affinities that exceed the canonical Hsp70 sequence (IEEVD) (**Fig 2a**). We anticipated that CHIC2, because it is expressed at substantially lower abundance than chaperones, might exploit a similar set of interactions to achieve tight binding, thus outcompeting the moderate affinity of the chaperones. Surprisingly, we found that the measured affinity of the labeled CHIC2 peptide (IFRPD, K_d_ 148 ± 6 nM) was not substantially tighter than that of Hsp70 (**Fig 2a**). Because the sequences of high affinity peptides such as CHIPOpt should be evolutionarily accessible, we wondered why CHIC2 retained a modest affinity sequence and how it might compete with abundant chaperones.

**Figure 2:**
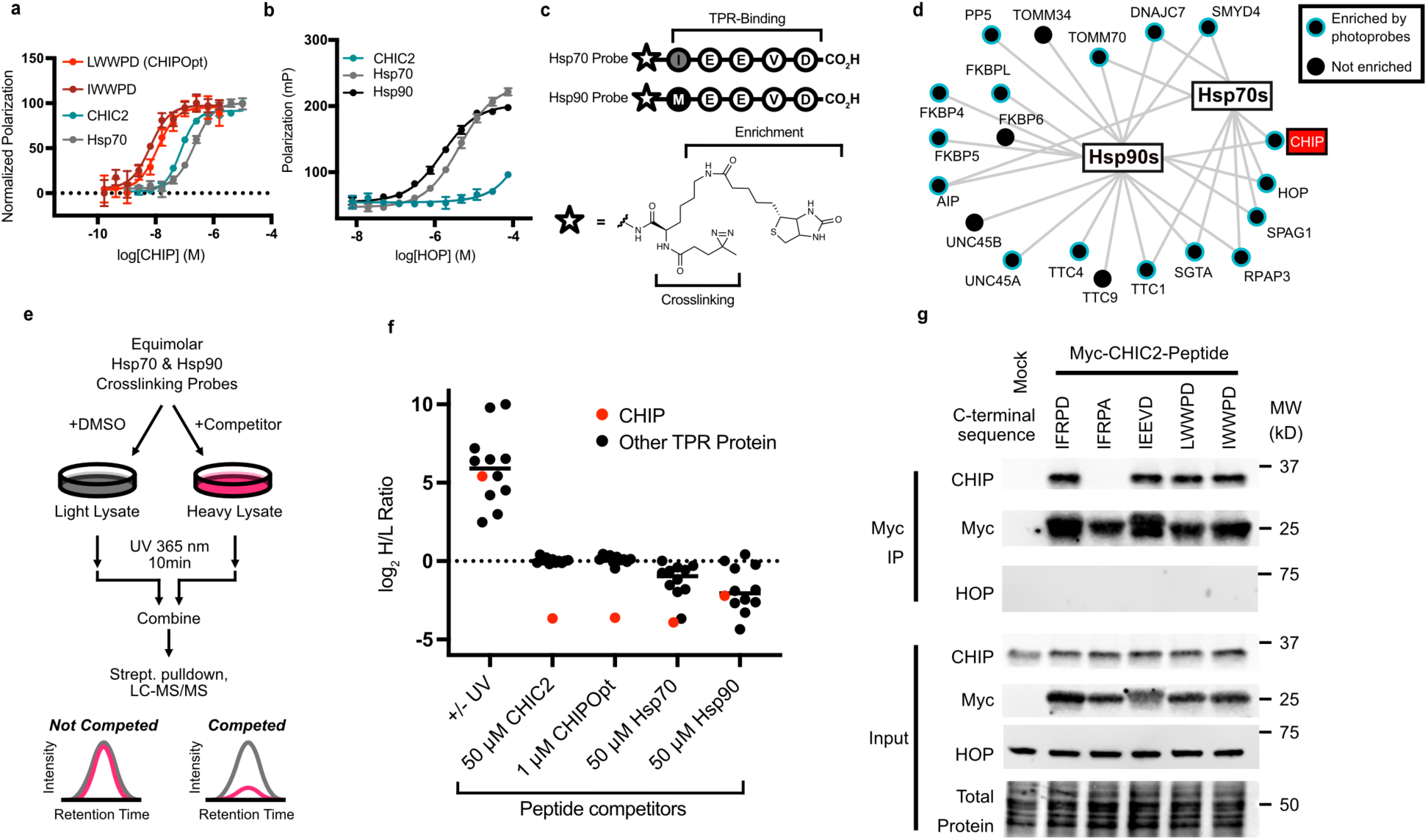
CHIC2 overcomes competitive chaperone binding to CHIP through selective interactions. **a)** Saturation binding of CHIP to fluorescence polarization tracers. Error bars represent S.D. of 4 replicate datapoints from 1 of 3 representative experiments. **b)** Saturation binding of fluorescent FP tracers to the related CC-TPR protein HOP. Error bars represent S.D. of 4 replicate datapoints from a single biological experiment. **c)** Design of photo-crosslinking probes for enrichment of TPR proteins. The C-terminal peptides of Hsp70 (IEEVD) and Hsp90 (MEEVD) were functionalized at their N-termini with an alkyl diazirine photocrosslinker and biotin enrichment handle, using a lysine residue as a spacer. **d)** Schematic map of known interactions (gray lines) between Hsp70 and Hsp90 and carboxylate clamp-containing TPR proteins, including CHIP. Proteins significantly enriched by either the Hsp70 or Hsp90 crosslinking probe (log2FC > 1 +/− UV treatment) in 293T cells are indicated in blue. **e)** Design of competitive chemoproteomics experiments for profiling peptide selectivity. Briefly, isotopically labeled lysates are incubated with crosslinking probes in the presence or absence of a peptide of interest, followed by crosslinking, streptavidin enrichment, and on-bead digest for subsequent LC-MS/MS analysis. Loss of signal in the heavy channel indicates substantial binding to a given TPR cochaperone of interest. **f)** Results of the competitive chemoproteomics experiment, showing the high selectivity of CHIC2 and CHIPOpt peptides. **g)** Co-immunoprecipitation of chimeric CHIC2 constructs from HEK293T cells. Substitution of the CHIC2 C-terminus with higher affinity peptides (LWWPD, IWWPD) did not confer additional CHIP binding, and fusion to the non-selective peptide IEEVD did not confer binding to the related TPR protein HOP. Results are representative of experiments performed in independent duplicates.

CHIP belongs to a family of ∼30 carboxylate-clamp containing TPR proteins (CC-TPR proteins) that are predicted to bind EEVD motifs^19^. Hsp70 and Hsp90 are known to bind many of these TPR proteins, and competition between them is thought to help regulate protein homeostasis^20^. Thus, we considered that an alternative way for CHIC2 to compete with Hsp70s and Hsp90s would be to enhance its selectivity for CHIP’s TPR domain over the other CC-TPR proteins. In other words, increased binding specificity might allow CHIC2 to effectively “seek out” free CHIP within the cell, while the chaperones are partitioned amongst the large number of other CC-TPR proteins (**Fig 2d**). As an initial test of this idea, we used FP to explore binding of C-termini from Hsp70, Hsp90, and CHIC2 to the relatively highly expressed CC-TPR protein HOP. While Hsp70 and Hsp90 had affinities for HOP that were in the expected, low micromolar range (Hsp70 K_d_ 4.4 µM, Hsp90 K_d_ 1.6 µM), the CHIC2 peptide was incapable of binding (**Fig 2b**). This result provided a first clue that CHIC2 might prefer CHIP over related CC-TPR proteins.

To test this hypothesis more broadly, we developed a chemical proteomics assay. Specifically, we coupled the IEEVD and MEEVD peptides from Hsp70 and Hsp90 to a biotin handle and diazirine photo-crosslinker (**Fig 2c**). We envisioned that, together, these probes might pull down the carboxylate clamp containing TPR proteins, including CHIP and HOP (**Fig 2d**). Using that system, we could then treat with the CHIC2-derived, C-terminal peptide and measure which interactions are inhibited using quantitative mass spectrometry (**Fig 2e**). After treatment of 293T cell lysates with a combination of the Hsp70- and Hsp90-derived probes and UV light, we confirmed that this chemoproteomics platform allows detection of up to 16 CC-TPR proteins in a given experiment (**Fig 2d, blue circles**). In key controls, we found that both probes showed UV-dependent enrichment and were effectively competed by unmodified versions of their parent peptides (**Extended Data 3**). Then, we treated the lysates with competitors and determined the extent to which each TPR protein was displaced. First, we confirmed that neither the Hsp70 nor Hsp90 peptides (50 µM) were highly selective; addition of these peptides displaced most of the CC-TPR proteins equally. In contrast, the designed ligand (CHIPOpt, 1 µM) was selective for CHIP as expected (**Fig 2f**). Strikingly, we found that the CHIC2 peptide (50 µM) was exquisitely selective for CHIP amongst CC-TPR proteins (**Fig 2f**), despite its modest affinity.

To explore whether this selectivity is inherent in the C-terminus of CHIC2, we generated chimeras of full length CHIC2 in which the native C-terminus (IFRPD) is replaced with the Hsp70 sequence (IEEVD) or the tighter binding designed peptides (LWWPD and IWWPD). We expressed the chimeras in 293T cells and measured binding to CHIP by co-IP. In those experiments, we found that deleting the terminal aspartic acid (IFRPA) blocked the interaction (**Fig. 2g**), as expected, and that the native and engineered sequences all bound similar levels of CHIP. However, fusion of IEEVD to CHIC2 did not reduce binding of the chimera to CHIP, nor did it confer binding to HOP, suggesting that interactions outside of CHIC2’s C-terminal motif must also contribute to selectivity. Thus, a combination of the IFRPD peptide’s intrinsic selectivity and PPIs outside the C-terminal region are responsible for mediating CHIC2’s selective interaction with CHIP in cells.

### CHIC2 binding promotes a dramatic reduction in CHIP ubiquitin ligase activity

Binding of CHIP to Hsp70’s EEVD motif leads to rapid polyubiquitination of Hsp70 *in vitro*^21^, so we expected a similar mechanism for the CHIC2 complex. To our surprise, *in vitro* ubiquitination experiments demonstrated that full length, human CHIC2 (see Methods) was exclusively and weakly mono-ubiquitinated (**Fig 3a-b, red stars**). This low level of ubiquitination might be expected if CHIC2 was somehow restricting CHIP’s enzyme activity, a possibility that can be conveniently measured by examining the auto-ubiquitination of CHIP. Indeed, when we increased the levels of CHIC2, the amount of CHIP auto-ubiquitination was drastically reduced (**Fig 3a-b**). To further test whether CHIC2 may inhibit CHIP’s ligase activity, and to rule out alternative effects on E1-mediated ubiquitin activation or E2 ubiquitin charging, we performed CHIP activity assays using a pre-charged E2 conjugate. Equimolar amounts of CHIC2 significantly reduced both E2 discharge and CHIP autoubiquitination in this context (**Fig 3c**), whereas super-stoichiometric amounts completely prevented discharge of the E2 conjugate. Therefore, CHIP’s ability to catalyze ubiquitin transfer seems to be dramatically reduced in the presence of CHIC2. To probe whether CHIC2 might also inhibit CHIP in the presence of a benchmark, physiological substrate, we added CHIC2 to reactions of CHIP with the microtubule associated-protein tau (MAPT/tau). Again, we found that CHIC2 largely blocked ubiquitin transfer to tau (**Fig 3d**). Importantly, adding a peptide derived from CHIC2’s EEVD-like motif was not able to fully replicate this effect (**Fig 3d**), suggesting that CHIC2’s full inhibitory activity required the intact protein. Taken together, these data support a model in which CHIC2 acts as a pseudosubstrate inhibitor of CHIP, using its EEVD-like motif to hinder recruitment of Hsp70 and, once fully bound, exploiting additional protein-protein interactions to reduce CHIP’s baseline catalytic activity.

**Figure 3:**
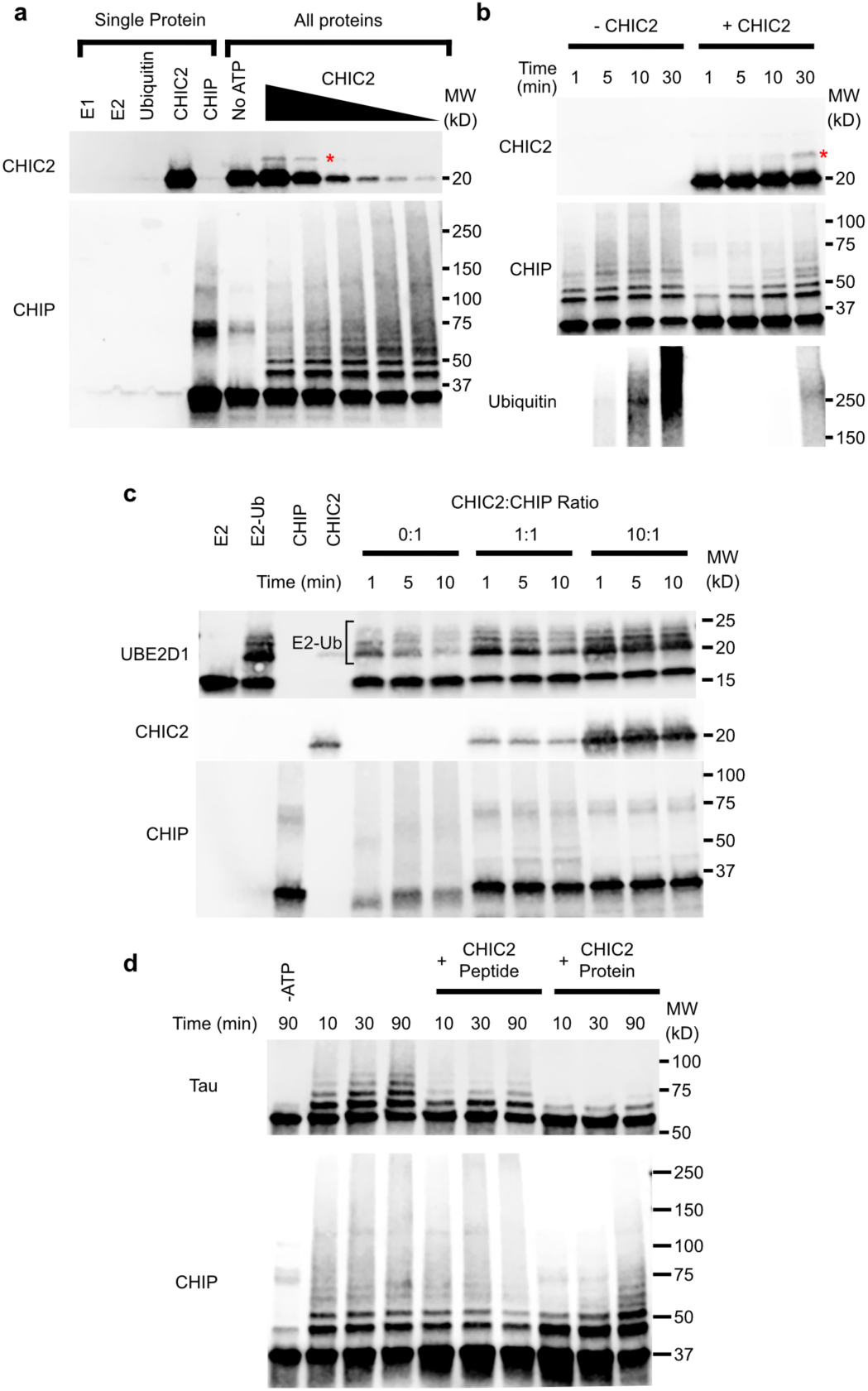
CHIC2 is monoubiquitinated and inhibits CHIP E3 ligase activity *in vitro*. **a)** *In vitro* ubiquitination reactions of CHIC2 (maximum 5 µM; serial 2-fold dilutions) and CHIP (0.5 µM; see Methods). Reactions were quenched after 10 minutes at RT. **b)** Time course of the CHIC2 (5 µM) ubiquitination reactions, performed as in panel a. **c)** Ubiquitination reactions using CHIP (0.5 µM) and pre-loaded UBE2D1-ubiquitin conjugate (0.5 µM; performed in the absence of ATP, ubiquitin or E1 enzyme; see Methods). Saturating concentrations of CHIC2 (10:1) inhibit CHIP’s ability to discharge E2 thioester conjugates. **d)** Ubiquitination of the known CHIP substrate tau is inhibited by CHIC2 protein (5 μM), but not CHIC2 peptide (50 μM). All results in this figure are representative of experiments performed in independent duplicates.

### Substrate recognition by CHIC2 and CHIP is highly coordinated in some cell types, but not others

We next wondered what effect CHIC2’s inhibitory activity might have on the proteome. CHIP and CHIC2 have been shown to regulate the levels of two members of the cytokine receptor family: interferon gamma receptor (IFNGR)^18,22,23^ and a subunit of multiple related cytokine receptors, CSF2RB^24^. These studies suggest that a CHIC2-CHIP complex may primarily regulate recycling and/or turnover of cytokine receptors. However, CHIP is also reported to regulate the levels of a variety of non-cytokine receptors, such as CFTR^25^, INSR^26^ and GHR^27^, so we wanted to more broadly explore the substrate scope of the CHIC2-CHIP complex. Accordingly, we generated CRISPR knockouts of either CHIC2 or CHIP in the glioblastoma line U251-MG (for characterization, see **Extended Data 4**) and then performed multiplexed quantitative proteomics to identify proteins that change abundance after deletion. We found that relatively few proteins were strongly regulated by knockout of either CHIC2 or CHIP (**Fig 4a**). Strikingly, however, a majority of the affected proteins were shared between the CHIC2 and CHIP knockouts (**Fig 4e**) and the fold-changes of these hits were also highly correlated (**Fig 4b**). This correlation was also true at the pathway level (**Fig 4c**), with two of the most impacted pathways being linked to cell surface receptor biology (**Fig 4d**). Other affected pathways included various intracellular responses, such as fatty acid metabolism, which might be due to direct or indirect effects of the knockouts. Importantly, these studies only explored changes in protein abundance, while the CHIC2-CHIP complex might also have effects on protein trafficking or other pathways. Nevertheless, the striking correlation between the proteins impacted by CHIC2 and CHIP knockout suggests that, in these cells (see below), a large majority of CHIP’s ability to regulate protein turnover seems to be mediated by the chaperone-independent, CHIC2 complex.

**Figure 4:**
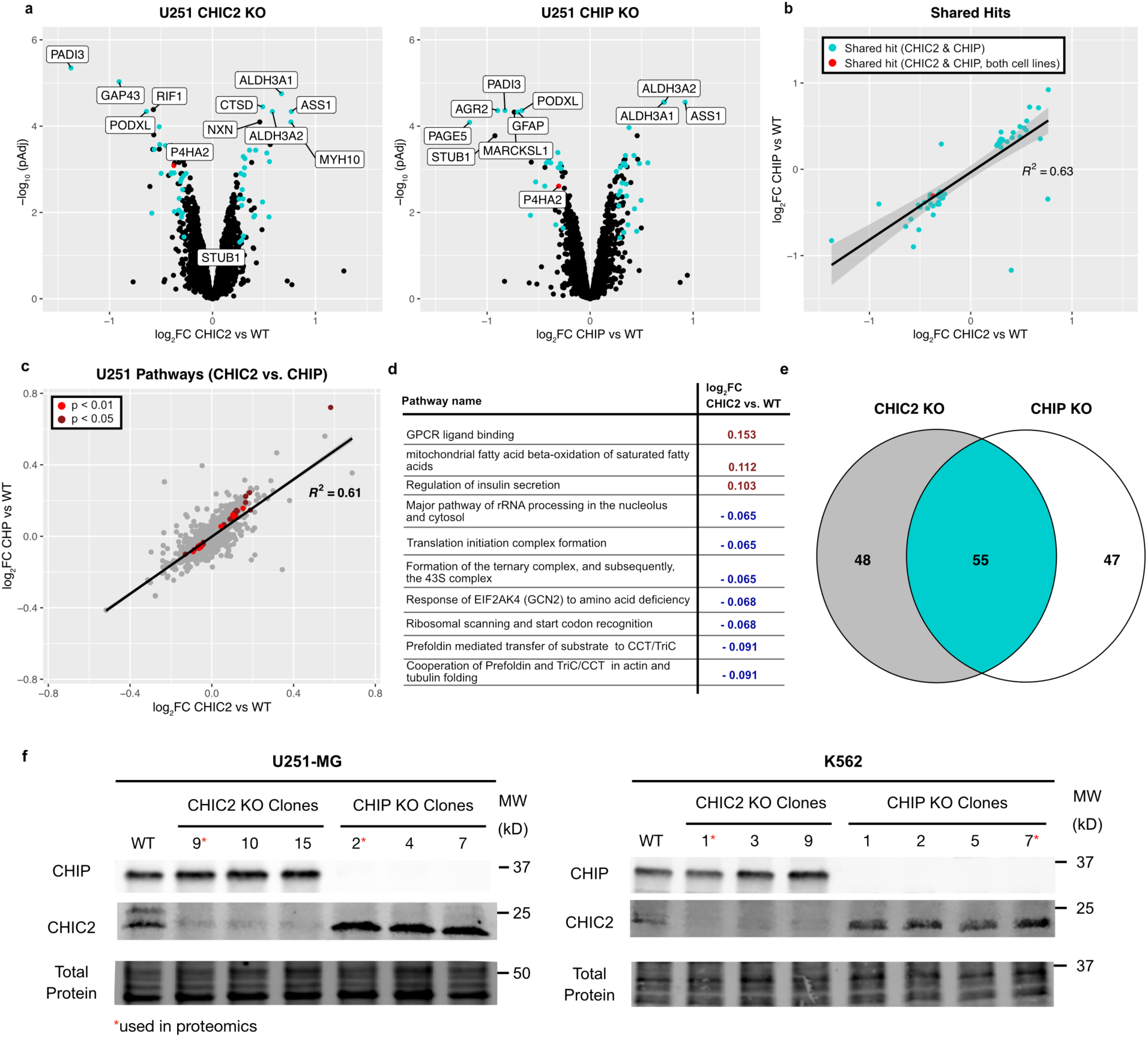
Substrate recognition by CHIC2 and CHIP is highly coordinated, but cell-type dependent. **a)** Results of TMT quantitative proteomics experiments, performed on parent U251-MG and CHIC2 and CHIP knockout U251-MG cells. Proteins with a log2FC >0.25 and a Benjamini-Hochberg adjusted *p*-value < 0.05 were considered hits. Blue dots indicate proteins whose abundance is significantly affected in both knockout conditions. **b)** Shared hits between CHIC2 and CHIP knockout have highly correlated fold-changes. Red dot indicates the one hit that is shared with the cell line K562 (see **Extended Data 5**). **c)** Analysis of pathways from the Reactome database that are affected by CHIC2 or CHIP knockout, using the correlation-adjusted mean rank gene set test (Camera)^44^. **d)** Top ten pathways (p < 0.01) identified as being shared between the CHIC2 and CHIP knockouts. **e)** Venn diagram of proteins that are differentially affected by CHIC2 and CHIP knockout (log2FC > 0.25, *p*Adj < 0.05). **f)** Western blots showing that CHIC2 levels are elevated by CHIP knockout in both U251 and K562 cells. At least three separate clones are shown for each knockout, with the red star indicating the clone used for proteomic experiments. See **Extended Data 4** for characterization of the clones. Results are representative of experiments performed in independent triplicates.

CHIP is highly expressed in the brain, and we found it compelling that U251-MG, a brain-derived cell line, exhibits chaperone-independent regulation through CHIC2. However, CHIP has been shown to work with chaperones in other contexts as well^28^. Therefore, to test whether cells from a different lineage might likewise display chaperone-independent regulation, we repeated the multiplexed quantitative proteomics experiments in a leukemia cell line, K562. In that cell type, we found a smaller amount of overlap between the proteins impacted by CHIC2 or CHIP knockout (**Extended Data 5**), resulting in a substantially weaker correlation between the fold-changes of CHIC2 and CHIP hits at the protein and pathway level. This effect was not due to a lack of interaction between CHIC2 and CHIP in K562 cells, as CHIC2 levels were reliably increased upon CHIP knockout in the same manner as in U251 cells (**Fig 4f)**. These results suggest that CHIP function depends less on the regulatory activity of CHIC2 in K562 cells, when compared to U251 cells. Another interesting observation from these comparative proteomics studies is that the responsive proteins and pathways were substantially different in the U251 and K562 cell lines (**Extended Data 5c-e**), consistent with CHIP’s role in regulating a broad, rather than narrow, set of substrates.

To further explore this cell type difference, we compared our proteomics datasets to those from previously reported CHIP knockout in melanoma cell lines^23^. While known CHIP substrates, such as IFNGR and JAK1, were some of the most significantly regulated substrates in these cell lines, they were substantially less significant hits in either U251 or K562 cells (**Extended Data 6a**) and the effect on IFNGR abundance was variable in experiments using K562 cells (**Extended Data 6b**). However, we noted that CHIC2 knockout partially stabilized IFNGR in both the U251 and K562 cell lines and increased its cell surface levels (**Extended Data 6c-e**), suggesting that some proportion of CHIP-mediated IFNGR regulation might typically depend on CHIC2. Together, these results highlight the complex nature of CHIP substrate regulation, and suggest that CHIC2 plays a key, and sometimes dominant, role in CHIP regulation depending on the cell type.

### Loss of the CHIC2 / CHIP interaction leads to decreased longevity & neurodegeneration in C. elegans

Our findings thus far suggest that chaperone-dependent and chaperone-independent mechanisms of CHIP function are both important, and that CHIC2 can be a dominant regulator of CHIP in some contexts. This model inspired us to consider the evolutionary origins of CHIC2. CHIC2 is a highly conserved protein across metazoans, and the C-terminal EEVD-like motif that is responsible for interacting with CHIP is also conserved (**Fig. 5a**). As an initial test as to whether the orthologs of CHIC2 and CHIP from another organism interact, we expressed CHIC (also known as TAG-266, the *C. elegans* ortholog of CHIC2) and CHN-1 (the *C. elegans* ortholog of CHIP) in 293T cells and confirmed that they bind in a manner that requires the same C-terminal aspartate (**Fig. 5b**).

**Figure 5:**
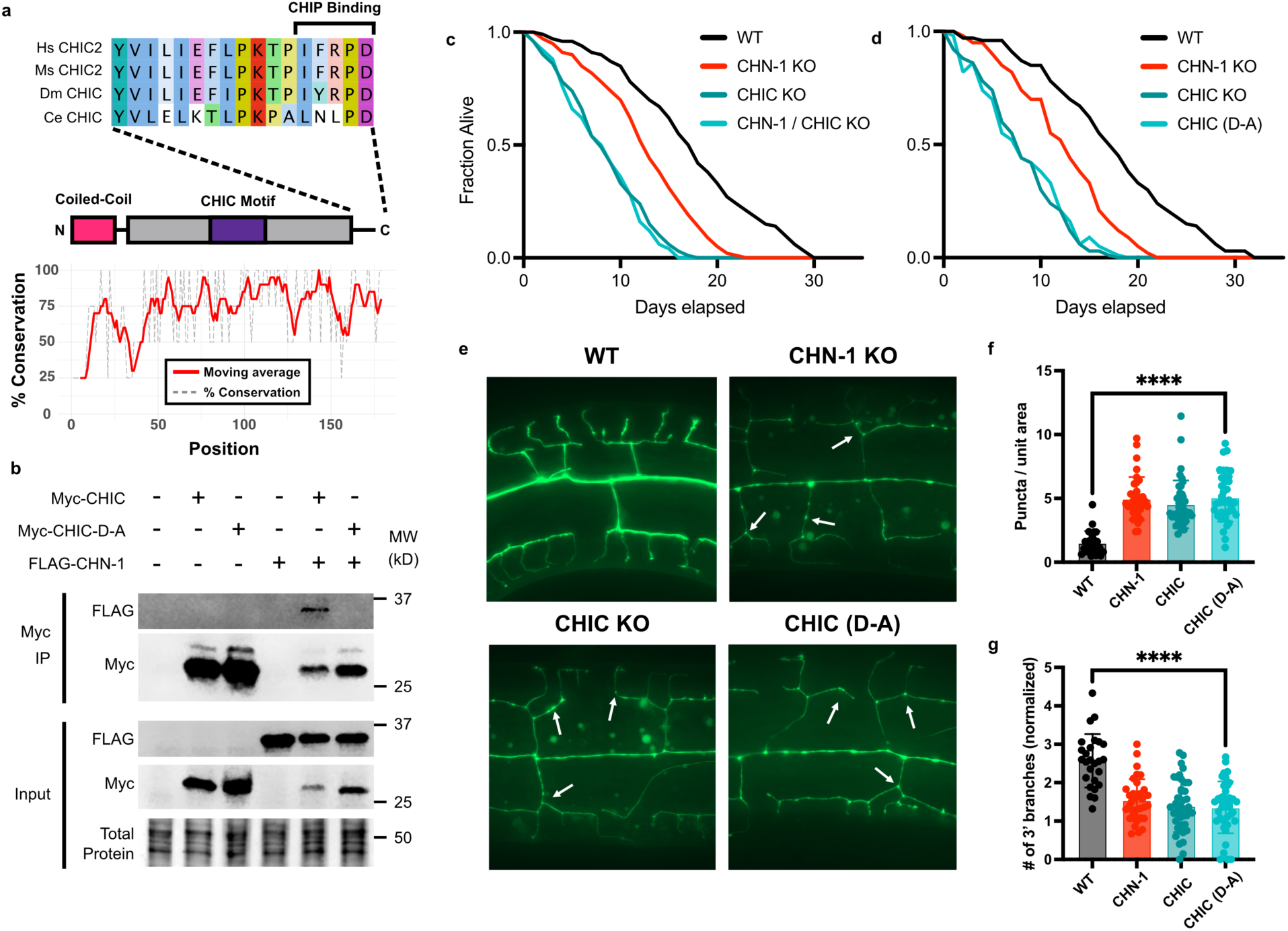
Loss of the CHIC2/CHIP interactions leads to decreased longevity and neurodegeneration in *C. elegans*. **a)** Sequence alignment of CHIC2 and CHIP orthologs from human, mouse, fruit fly, and worm illustrate a high degree of conservation, including in the C-terminal EEVD-like motif that binds CHIP (upper inset). **b)** Results of co-immunoprecipitation experiments from HEK393T cells expressing *C. elegans* myc*-*CHIC (ortholog of CHIC2) and FLAG-CHN-1 (ortholog of CHIP). Data is representative of 2 independent biological experiments. **c)** Results of longevity assays in *C. elegans*. Data represent the average of 3 independent experiments using 75 animals each. **d)** Results of longevity assays with a CRISPR-edited strain in which CHIC’s C-terminal Asp is replaced with Ala (CHIC D-A), showing that loss of the CHIC/CHN-1 interaction decreases longevity. **e)** Knockout and mutant strains display increased rates of PVD neuron degeneration in the strain *wdls51,* which selectively expresses GFP in this cell type. Formation of bead-like puncta on the PVD dendrite (arrows) is indicative of neurodegeneration. **f)** Quantification of neurodegenerative puncta as in panel e). Puncta were counted by a blinded observer, and then normalized to the total area of each image. At least 25 animals per strain were quantified, and data are representative of 3 independent biological experiments. **** *p* < 0.0001 by Welch’s *t*-test. **g)** Loss of branched PVD neuron density in mutant strains. Dendrite branches were quantified by a blinded observer who counted junctions moving outward from the central axon. The number of 3’ branches was normalized to the number of parental (1’, 2’) branches. At least 25 animals per strain were quantified, and data are representative of 3 independent biological experiments. **** *p* < 0.0001 by Welch’s *t*-test.

*C. elegans* is an ideal model for further studying the biological roles of this complex because both CHIP and its orthologue CHN-1 have been shown to regulate longevity^26,28^. Thus, we hypothesized that disruption of the CHIC-CHN-1 interaction might phenocopy this effect, but only if much of CHN-1’s activity is mediated through CHIC. As an initial test of this idea, we knocked out CHIC and found that surprisingly, it had a substantially more detrimental effect on worm longevity than CHN-1 knockout (**Fig 5c**). Given our observation that CHIC2 inhibits CHIP ligase activity (see Fig. 3), we reasoned that toxic gain-of-function upon removal of CHIC inhibitory activity might contribute to this strong phenotype. However, the CHN-1/CHIC double knockout did not restore viability to the level of the CHN-1 knockout, suggesting that CHIC may have additional functions beyond simply attenuating CHN-1 activity. In order to exclude potential CHN-1-independent impacts of CHIC knockout, we next used CRISPR editing to replace the endogenous CHIC gene with a mutant lacking the terminal aspartate residue (CHIC D-A). This edit phenocopied CHIC knockout (**Fig. 5c**), rigorously confirming that the interaction with CHN-1 is essential for proper CHIC function. Taken together, these results show that the chaperone-independent interaction between CHIC and CHN-1 is essential for longevity in an animal.

As in mammals, CHIC and CHN-1 are highly expressed in the *C. elegans* nervous system^29,30^. Thus, we were curious whether disruption of the CHIC/CHN-1 interaction might be sufficient to impact neuronal health. Using an imaging assay^31^, we found that CHN-1 and CHIC knockout worms exhibited abnormal neuronal branching and increased puncta in PVD neurons (**Fig. 5e**, arrows, & **Fig 5f**), consistent with impaired neuronal health and neurodegeneration. The knockout strains also showed significantly less peripheral neuronal development (**Fig 5g**). Again, even mutation of a single amino acid in the C-terminus of CHIC (CHIC D-A) was sufficient to trigger these phenotypes. Together, these results validate the importance of CHIC2’s interaction with CHIP in an organismal context. The stringent conservation of this interaction and its dramatic impact on longevity and neuronal health support a fundamental biological role for chaperone-independent regulation of CHIP in metazoans.

## Discussion

CHIP has been implicated in a wide variety of biological processes^32^. For example, mutations in CHIP are linked to severe neurodegenerative disease in humans^33,34^ and the CHIP^−/−^ mouse develops inclusions of tau^35^. It has previously been assumed that these functions are largely a product of CHIP working in concert with the molecular chaperones, Hsp70 and Hsp90^36^. In that canonical mode, the chaperones bind to misfolded proteins and recruit CHIP with their EEVD motifs, favoring polyubiquitination and turnover. Here, we have described an alternative interaction with the membrane-anchored protein, CHIC2, that competes with chaperone binding and restricts CHIP function. Surprisingly, we found that this chaperone-independent role can sometimes predominate, as the proteomic changes induced by CHIC2 and CHIP knockout can largely phenocopy one another in U251 cells (see Fig 4). Furthermore, mutating a single aspartate that blocks binding to the CHIC2 ortholog, CHIC, induces premature aging and neurodegeneration in *C. elegans* (see Fig 5). These results demonstrate a significant and previously underappreciated role for CHIC2 in shaping CHIP’s overall biological functions.

Hsp70s and Hsp90s are highly abundant proteins, reaching ∼1% of soluble protein content in some cells^37^. At first glance, it seems challenging for CHIC2 to compete amongst this large pool of EEVD motifs, especially because its affinity is not substantially different than those of the chaperones (see Fig 2). Indeed, in K562 cells, the effects of CHIP and CHIC2 knockout were somewhat decoupled (see Extended Data 5), suggesting that, broadly, CHIP does prefer to work with other partners such as chaperones in that cell line. How then do we harmonize these contrasting examples of chaperone-dependent and chaperone-independent CHIP function? Our chemoproteomics results (see Fig 2) provide a key insight here, because CHIC2 seems to use selectivity, rather than outright affinity, as one way to tip the balance in its favor. As a framework, we envision a model in which the relative expression of CHIP and its partners dictates whether chaperone-dependent or chaperone-independent mechanisms will dominate (**Figure 6).** Briefly, while chaperones must interact with a large pool of other CC-TPR proteins, CHIC2 has an EEVD-like motif that is selective for CHIP. Thus, formation of the CHIC2/CHIP complex is only sensitive to the amount of free CHIP, whereas formation of a chaperone-bound CHIP complex is dependent on both the relative concentrations of the chaperones and their many other binding partners. This distinct selectivity may also allow cells to regulate the behavior of CHIP independently from related CC-TPR proteins.

**Figure 6:**
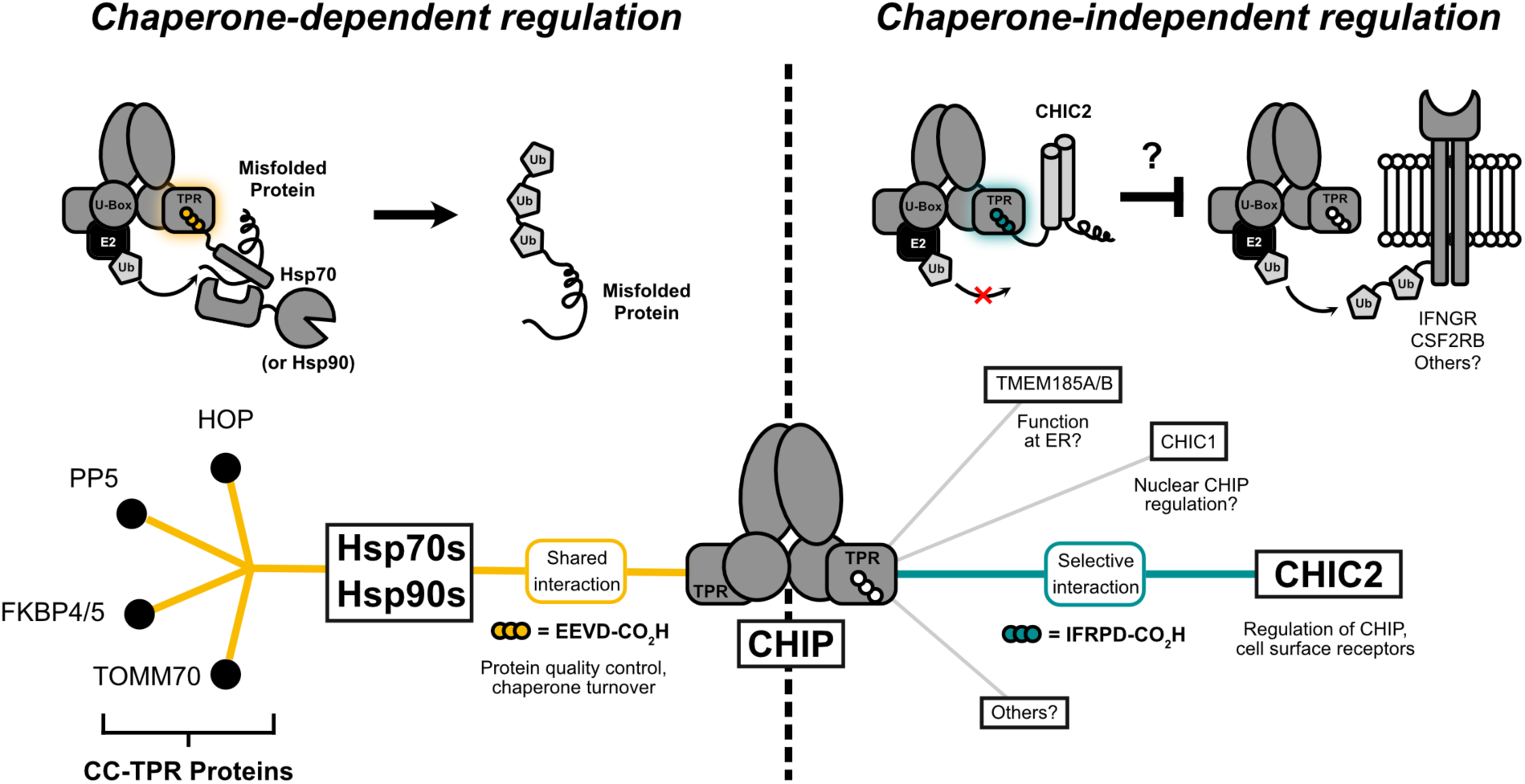
A model for context-specific regulation of CHIP. Competition between chaperone-dependent (left) and chaperone-independent (right) interactions dictate CHIP behavior in different contexts. Binding of “free” CHIP (center) to various partners is governed by the relative expression levels of Hsp70s & Hsp90s, CC-TPR proteins, and chaperone-independent interactors such as CHIC2. Since CHIC2 interacts selectively with CHIP, its interaction is influenced by free CHIP levels but not that of numerous other CC-TPR proteins. Relative levels of free chaperone and CHIP may therefore determine the “set point” at which formation of the CHIC2/CHIP complex occurs in different cell types. Other factors such as PTMs, chaperone upregulation under stress, or subcellular localization may further tune this behavior. This complex web of interactions likely drives the cell-type specificity of CHIP behavior and its response to various stressors.

Within this model, it is interesting to speculate that the ability of CHIC2 to interact with CHIP might be further tuned under specific conditions. For example, the levels of Hsp70 and Hsp90 are highly elevated by heat shock, and CHIP is known to regulate their return to basal expression following resolution of stress^7^. Thus, CHIP may be transiently displaced from CHIC2 under such conditions. Conversely, phosphorylation of Hsp70’s EEVD motif is known to block its interaction with CHIP, but not other CC-TPR proteins^38^, which might transiently free CHIP to bind CHIC2. Given the link between CHIC2 and cell surface receptor regulation, this may be a way for cells to change how they respond to external stimuli (*e.g.* cytokines^22,24^) when internal stress pathways are activated. In other words, the relative expression levels of CHIC2, CHIP, and chaperones might dictate the “set point” at which the transition from chaperone-dependent to chaperone-independent CHIP function occurs across cell types. This model could partially explain the cell-type specificity that we observed in our proteomics experiments. While we are far from fully modelling this complex network of PPIs, we propose that the balance of relative affinities, PTMs and protein concentrations seems likely to dictate the partners of CHIP, shaping its substrates, localization, and function in dynamic ways.

More broadly, this work also has implications for CHIP’s recognition of other chaperone-independent substrates beyond CHIC2. We note that CHIPScore predicts a number of other proteins to interact strongly with CHIP (**Extended Data 8**). For example, CHIC1 is highly homologous to CHIC2 and is able to recruit CHIP through the same EEVD-like motif (**Extended Data 8a, c**). It therefore seems likely that CHIC1 would also bind and inhibit CHIP. However, there is no evidence to link CHIC1 and CHIP in the DepMap, which implies that CHIC1 and CHIC2 do not perform redundant functions. Surprisingly, we found that CHIC1 is predominantly localized to the nucleus, not the membrane (**Extended Data 8b**). Therefore, CHIC1’s role might be to restrict CHIP function in a distinct subcellular location. Two other predicted CHIP partners, TMEM185A and TMEM185B, are 7-transmembrane proteins of unknown function that are expressed in the endoplasmic reticulum (ER)^39^, and their protein levels increase significantly upon CHIP knockout in various cell types (**Extended Data 8e).** Co-IP studies in 293T cells showed that these proteins form a chaperone-independent complex with CHIP, but that CHIC2 was excluded (**Extended Data 8d**). Therefore, TMEM185A/B might assemble a distinct complex with CHIP at the ER. Formation of such complexes may be further modulated by response to stress stimuli, as previous work has revealed hundreds of additional EEVD-like motifs that are generated by proteolysis^11^. We anticipate that additional, important CHIP partners remain to be discovered and characterized.

Because of its subcellular localization, it seems likely that CHIC2 would predominantly shape CHIP’s behavior at the membrane (**Figure 6**). While it is not yet clear what exact mechanisms are at play, there are several compelling possibilities that deserve future consideration. Given our *in vitro* ubiquitination results (see Fig 3), monoubiquitination of CHIC2 seem likely to occur in this environment. Indeed, a band consistent with the molecular weight of monoubiquitinated CHIC2 disappears upon CHIP knockout in U251 cells, despite total CHIC2 levels increasing (see Fig 4f). Monoubiquitination has been strongly linked to vesicular transport^40^, which may serve to modulate endocytic trafficking of receptors such as CSF2RB^24^. Thus, monoubiquitination of CHIC2, and potentially nearby substrates, could be involved in regulation of receptor trafficking. Mono-ubiquitination of CHIP itself has also been recently shown to change its oligomeric state and processivity^41^, so modulation of CHIP’s autoubiquitination activity by CHIC2 might also be a contributing factor. Finally, it is also possible that CHIC2 may simply serve to inhibit CHIP’s chaperone-dependent activity in specific, membrane-proximal regions, thus protecting some substrates from degradation. We propose that key clues to these possible mechanisms might emerge by studying the dysfunction of CHIC2 in disease. Deletion or fusion of CHIC2 is observed in certain leukemias and systemic mastocytosis^42,43^, and copy number variations at the CHIC2 locus are common across multiple cohorts in The Cancer Genome Atlas. Therefore, studying these disease systems may uncover key substrates and mechanisms of the CHIC2/CHIP complex.

## Acknowledgements

This work was supported by grants from the NIH (RF1AG068125 to J.E.G.; F31AG077842 to M.D.C., RF1NS127414 to A.W.K., and R35GM128595 to R.C.P.), and by a National Science Foundation Graduate Research Fellowship to M.D.C. The results shown here are in part based on data generated by the TCGA Research Network: https://www.cancer.gov/tcga. Molecular graphics and analyses were performed with UCSF Chimera, developed by the Resource for Biocomputing, Visualization, and Informatics at the University of California, San Francisco, with support from NIH P41-GM103311.

We would like to thank Arun Wiita and Emilio Ramos for their assistance in conducting and interpreting our proteomics experiments, Martin Kampmann for his assistance in generating CRISPR knockouts, and Eric Greene for his helpful feedback on the manuscript.

## Author Contributions

M.D.C. conceived the study, designed, and performed experiments. M.H. designed and performed *C. elegans* experiments. E.C.C. and C.M.N. aided in protein expression / purification and helped design experiments. M.R. designed and helped develop the chemical proteomics assay. A.R.I.S., A.R.C., L.E.M., J.C.N., and R.C.P. performed X-ray crystallography and structure refinement. M.C. and J.G. wrote the manuscript, and all authors approved the final version.

---

**Extended Data 1:**
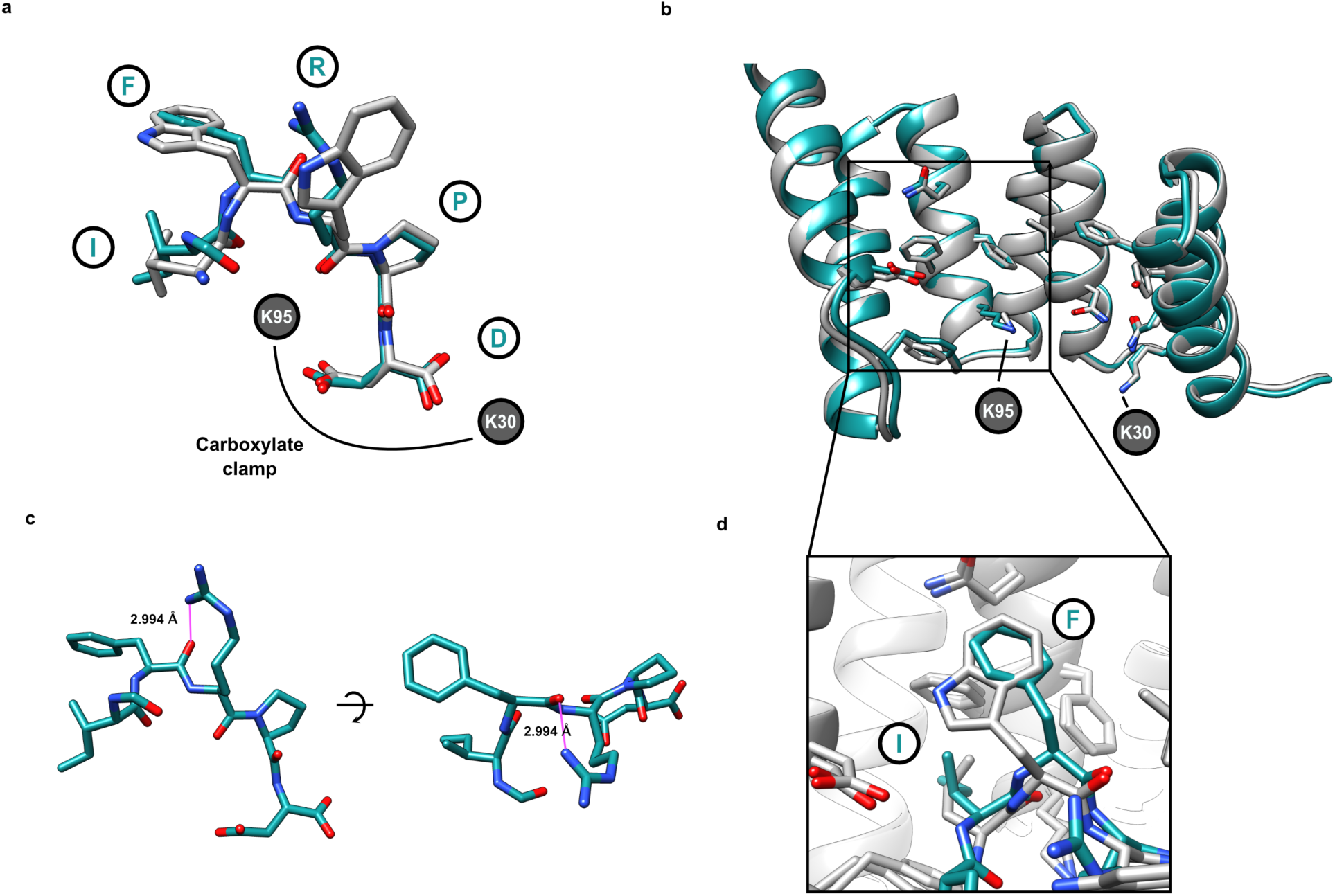
CHIC2’s EEVD-like motif bound to CHIP’s TPR domain adopts a similar conformation as the benchmark CHIPOpt. **a)** Overlay of the CHIP-bound conformation of the last five amino acids of CHIC2 peptide (IFRPD, cyan) and CHIPOpt (LWWPD, gray) demonstrates a similar binding pose. **b)** Comparison of CHIP’s TPR domains bound to either peptide (see panel a) also suggests a similar structure. **c)** An internal hydrogen bond between the P3 arginine side chain and the peptide backbone in the CHIC2 peptide may help pre-organize or stabilize the CHIP-binding mode. **d)** Similar contacts define the “hydrophobic shelf” in CHIP’s TPR domain and its interactions with the P5 isoleucine and P4 phenylalanine in CHIC2, compared to the P4 valine and P5 tryptophan in CHIPOpt. We speculate that the slight differences in backbone conformation may be enforced by the internal hydrogen bond highlighted in panel c).

**Extended Data 2:**
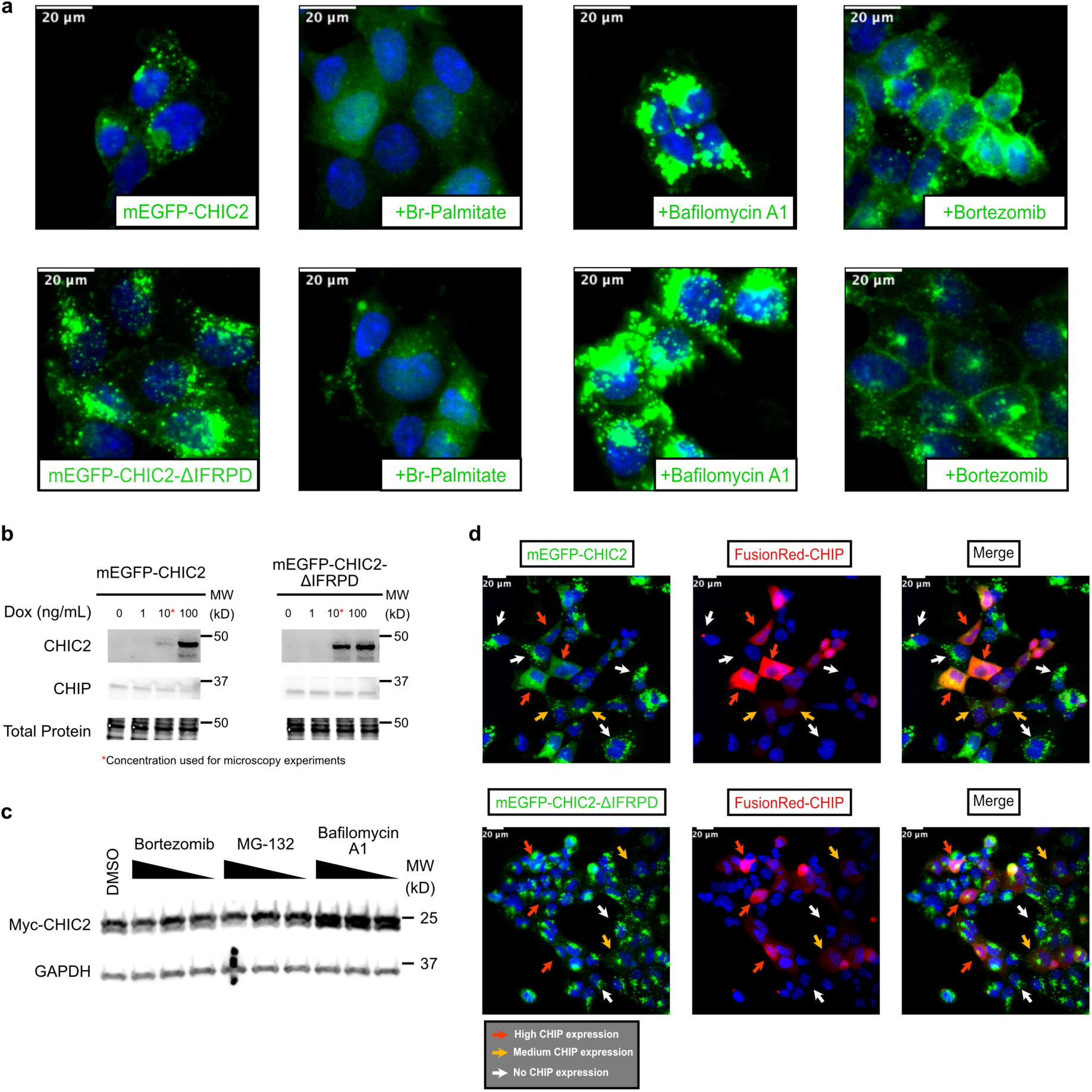
Subcellular localization of CHIC2 in response to inhibitor treatment and CHIP overexpression. **a)** Live-cell imaging of dox-inducible HEK293 Flp-In cells expressing either GFP-CHIC2-WT or GFP-CHIC2-ΔIFRPD. The C-terminal mutant exhibits higher basal expression levels of CHIC2, suggesting that CHIP is involved in CHIC2 clearance. Bafilomycin treatment induces a substantial accumulation of both WT CHIC2 and CHIC2-ΔIFRPD relative to bortezomib, suggesting that the lysosome is predominantly responsible for clearance, regardless of CHIP’s involvement. **b)** Western blots showing GFP-CHIC2 levels in response to doxycycline titration. The minimal amount of doxycycline that induced observable expression (10 ng/mL) was used in microscopy experiments. **c)** Western blots showing the results of transfection of HEK293T cells with Myc-CHIC2, followed by a 4 hour treatment with proteasome (bortezomib or MG-132) and lysosome (bafilomycin) inhibitors, suggesting that CHIC2 is predominantly degraded in the lysosome. Compounds were serially diluted 1:10 from 10 μM (bortezomib, MG132) or 1 μM (bafilomycin A1). **d)** Live cell microscopy, showing that high levels of FusionRed-CHIP overexpression (red arrows) result in a substantial relocalization of CHIC2-WT, but not ΔIFRPD. Intermediate expression of FusionRed-CHIP (yellow arrows) are not sufficient, suggesting that CHIP expression must reach a critical level to relocalize CHIC2. To limit the impact of degradation, cells were treated with bafilomycin (100 nM) for 4 hours prior to imaging.

**Extended Data 3:**
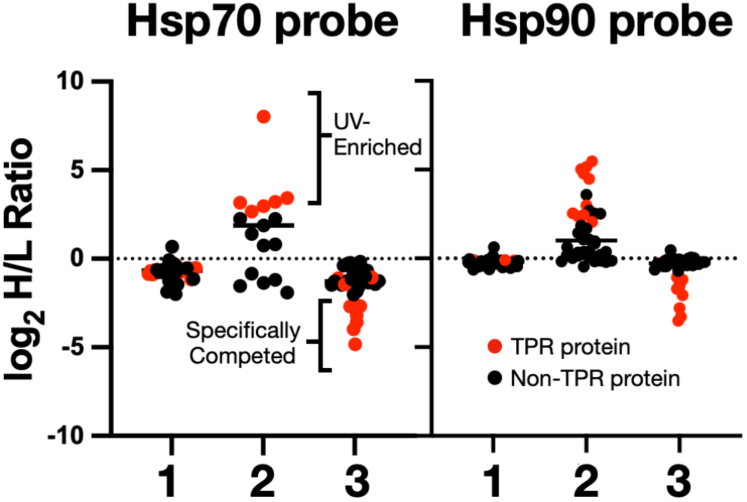
TPR protein enrichment by Hsp70 and Hsp90 photoprobes. **1)** Equal concentrations of Hsp70 or Hsp90 probes (10 μM; see Fig 2c) were added to both light and heavy channels, demonstrating balanced incorporation of isotopic labels. **2)** Results from experiments in which either Hsp70 or Hsp90 probe (10 µM) was added to each channel, but the light channel was not exposed to UV light, demonstrating specific crosslinking and enrichment. **3)** Hsp70 or Hsp90 peptide competitor (100 µM) was added to the heavy channel prior to crosslinking. For all experiments, H/L ratios were directly exported from MaxQuant.

**Extended Data 4:**
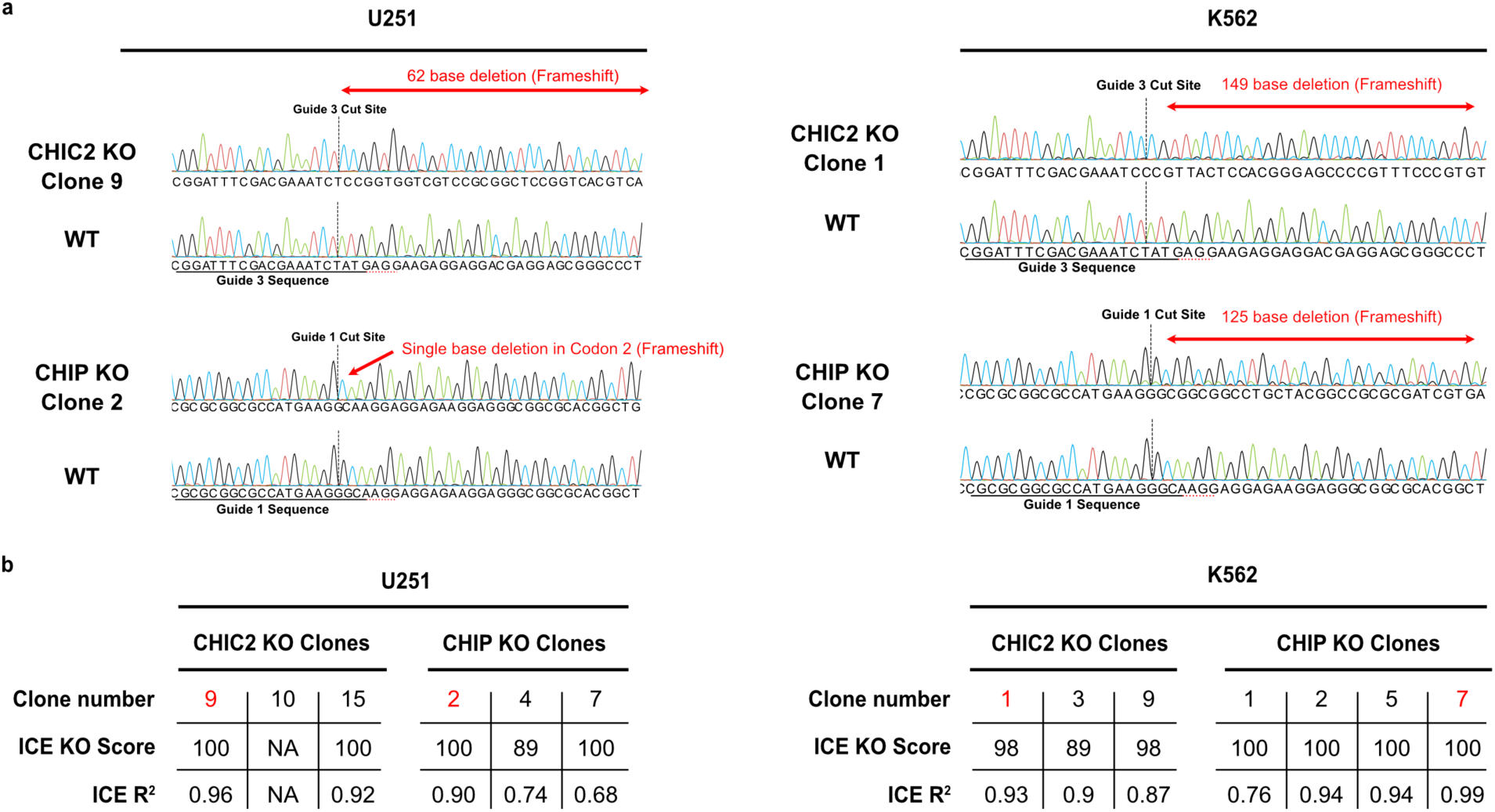
Characterization of CHIC2 and CHIP knockout clones. **a)** Sanger traces of selected knockout clones highlighting indels around the CRISPR cut sites. Guide sequences are underlined in the WT sequence. **b)** Summary of Sanger sequencing analysis for all clones using the Synthego ICE analysis software. NA indicates that ICE failed to generate a score from the sequencing files.

**Extended data 5:**
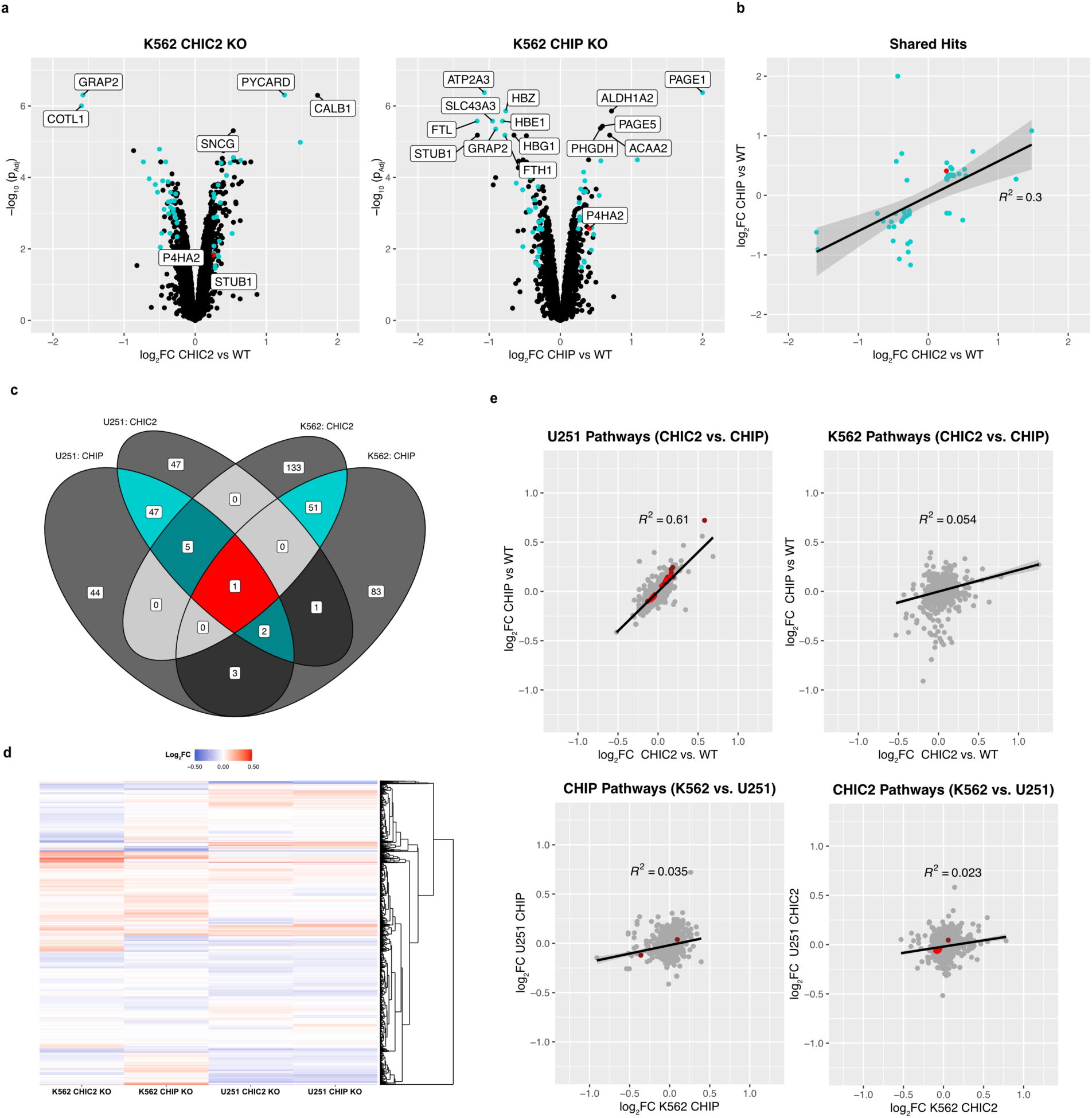
Proteomic changes in response to CHIC2 and CHIP knockout in K562 cells, highlighting cell type differences. **a)** Results of TMT quantitative proteomics of CHIC2 and CHIP knockout in K562 cells. Proteins with a log2FC >0.25 and a Benjamini-Hochberg adjusted *p*-value < 0.05 were considered hits. Blue dots indicate proteins for which the change in abundance is shared between the two knockouts. **b)** Plot of the fold-change in protein abundance, comparing the CHIC2 and CHIP knockout and showing the poor correlation between the shared hits. Some hits are elevated in abundance in one knockout and reduced in the other. Red dot indicates the one hit that is shared with the U251 cell line (see Figure 4). **c)** Venn diagram of hits shared between knockout conditions in the U251 and K562 cell lines, showing that the majority of overlapping hits are shared within a cell line rather than across a cell line or gene. **d)** Heatmap of proteomic fold-changes (KO vs. WT). Hierarchical clustering reveals a strong relationship within, but not across, cell lines. **e)** Pathway analysis from the Reactome database, using the correlation-adjusted mean rank gene set test (Camera)^44^. Pathways that are differentially regulated in response to CHIC2 or CHIP knockout are not shared between cell lines.

**Extended Data 6:**
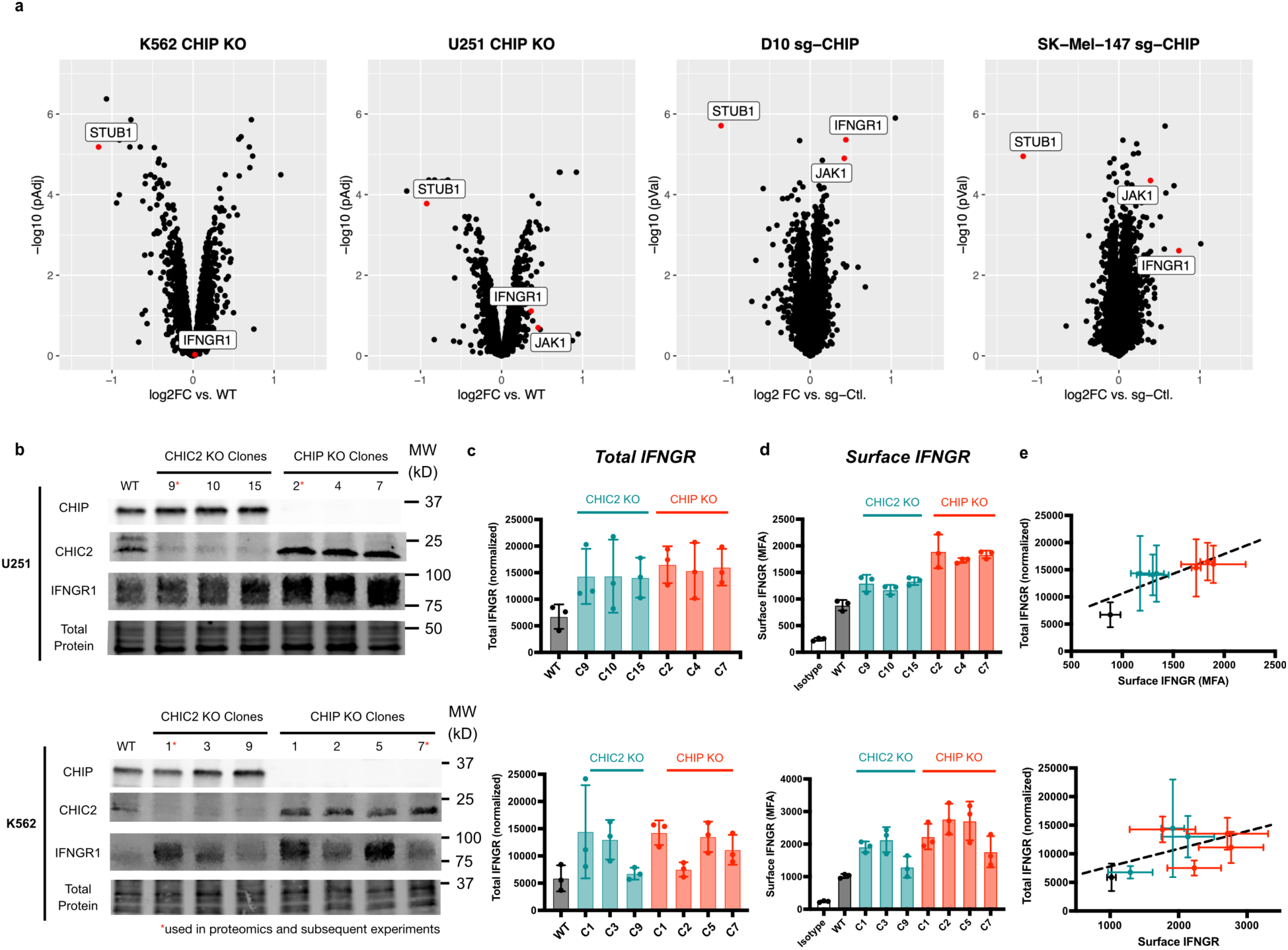
Cell-type dependent regulation of IFNGR by CHIP and CHIC2. **a)** Comparison of quantitative proteomics datasets from different CHIP knockout cell lines, highlighting the cell-type-dependent regulation of IFNGR. The results from the D10 and SK-Mel-147 cell lines is re-plotted from Apriamashvili et al^23^. Note that IFNGR1 and JAK1 were only identified by a single peptide in the K562 and U251 cell lines. **b)** Western blots of U251 and K562 knockout clones, showing that IFNGR upregulation in response to CHIC2 or CHIP knockout is consistent across clones in U251 cells, but more variable in K562 cells. In contrast, CHIC2 levels are consistently elevated upon CHIP knockout. **c)** Quantification of three independent western blotting experiments as in b). Error bars represent the S.D. of all the experiments. **d)** Flow cytometry analysis of surface IFNGR upon CHIC2 and CHIP knockout, showing increases in IFNGR surface levels. Error bars represent the S.D. of three independent biological experiments. **e)** Correlation of total and surface levels of IFNGR in response to knockout of CHIC2 (blue dots) and CHIP (red dots) across various clones, highlighting the strong correlation across clones in U251, but not K562. Error bars represent the S.D. of all the experiments.

**Extended data 7:**
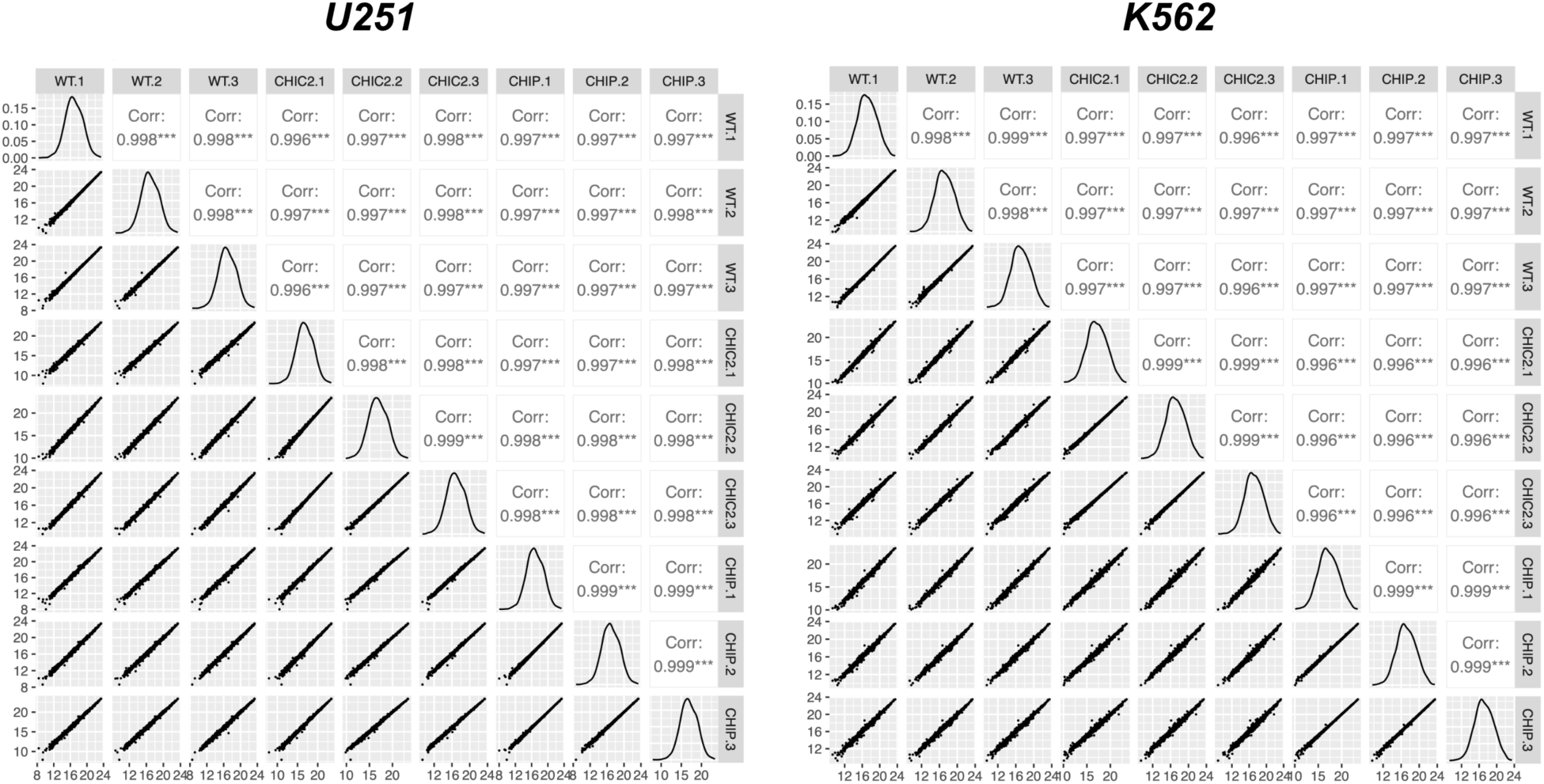
The TMT proteomics experiments show high replicate reproducibility. Median normalized protein intensities are highly reproducible across biological replicates.

**Extended Data 8:**
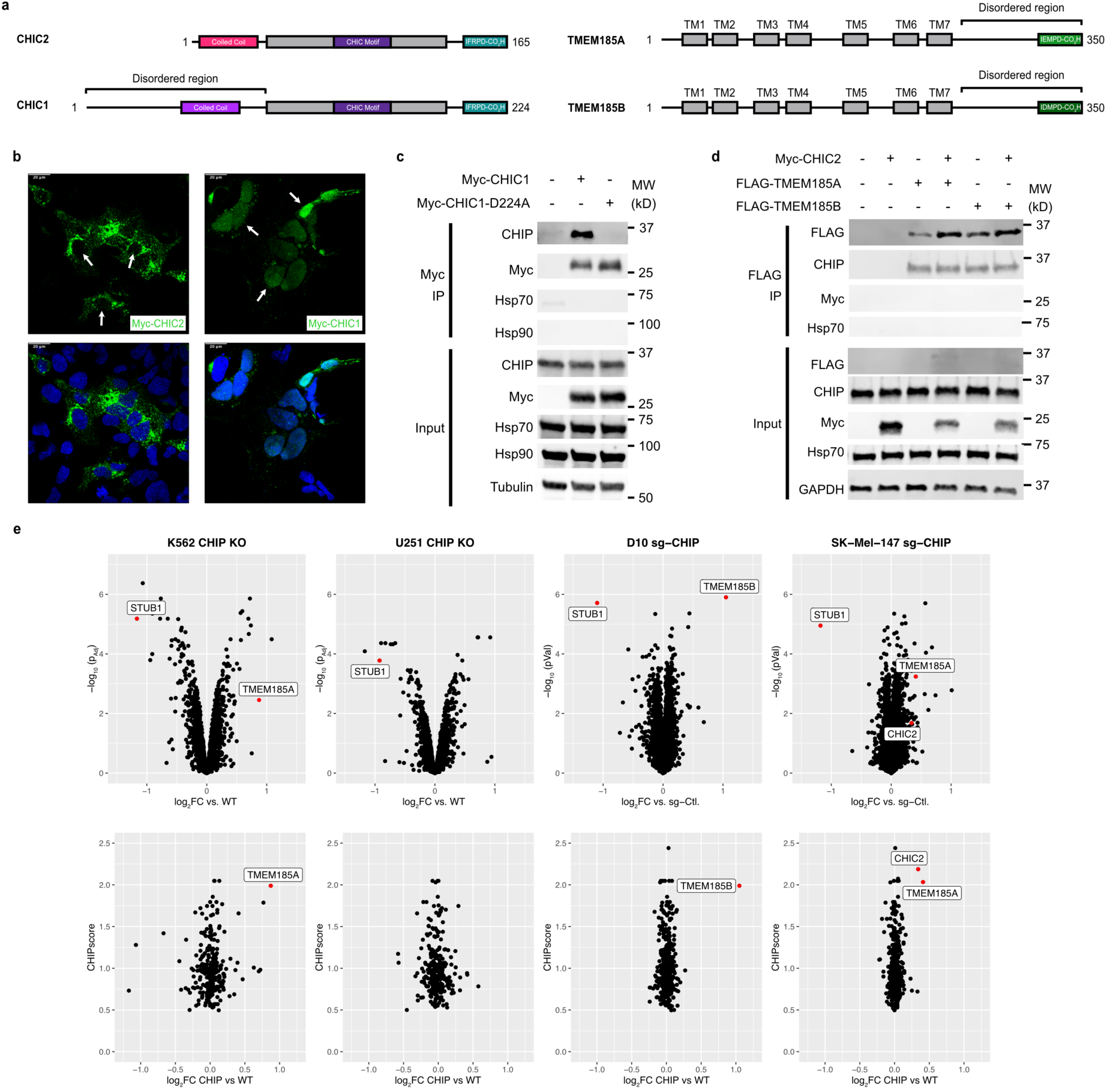
Chaperone-independent interactors of CHIP beyond CHIC2. **a)** CHIPScore also returns additional proteins with putative CHIP-binding EEVD-like motifs. The domain architecture of three of these factors: CHIC1, TMEM185A, and TMEM185B, are shown, with the putative EEVD-like motif highlighted and compared to CHIC2. **b)** Immunofluorescence microscopy results demonstrating nuclear localization of myc-CHIC1 (arrows) in HEK293 cells, compared to the primarily vesicular localization of myc-CHIC2. Myc-CHIC2 seems excluded from the nucleus (arrows). Scale bars (upper left) are 20 μm. **c)** Co-immunoprecipitation of myc-CHIC1 and CHIP in HEK293T cells demonstrates a dependence on the equivalent C-terminal aspartate as in CHIC2. Results are representative of independent experiments performed in duplicate. **d)** Co-immunoprecipitation of TMEM185A and TMEM185B from HEK293T cells shows an interaction with CHIP, but that CHIC2 is excluded. Moreover, over-expression of CHIC2 does not seem to interfere with the TMEM185A/B interaction with CHIP. Together, these results suggest that the TMEM185A/B and CHIC2 complexes with CHIP are distinct. **e)** Quantitative proteomics demonstrates that the levels of TMEM185A/B are consistently elevated in response to CHIP knockout across cell lines. Similar to what is observed for CHIC2, plotting the fold-change of TMEM185A/B compared to their CHIPScores (lower panels) suggests that these chaperone-independent CHIP complexes might be biologically important. Note that TMEM185A was only identified by a single peptide in K562 cells, and was not identified in U251.

## Methods

### DepMap analysis

The top 100 co-dependencies of CHIP (by CRISPR) were downloaded from the Cancer Dependency Map (22Q1 release), and the CHIPScore calculated for each. The top 100 CHIP-scoring proteins in the proteome were also selected (independent of DepMep association), and their genetic codependency with CHIP was also extracted. Proteins without a C-terminal aspartic acid were excluded from the analysis, as were genes not screened by CRISPR in the DepMap.

### Fluorescence polarization

#### Saturation binding

CHIP was serially diluted to 2X its final concentration in assay buffer (25 mM HEPES pH 7.4, 50 mM KCl, 0.01% Triton X-100, 1mM TCEP). Protein dilutions were then mixed 1:1 with a 2X tracer (FAM-Ahx-Peptide) solution in assay buffer + 2% DMSO to yield a final concentration of 1 nM tracer + 1% DMSO. From the resulting dilutions, an aliquot (18 μL) was pipetted into black, low-volume 384-well plates (Corning 4511) and incubated at RT for 15 minutes. Fluorescence polarization was measured on a BioTek H4 plate reader. Raw polarization data (mP) was normalized to buffer controls, then plotted relative to log_10_[Protein]. Data was fit to the log[agonist] vs. response model with variable slope in GraphPad PRISM 9.0, and K_d_ values were extrapolated from half-maximal effective concentration (EC_50_). HOP saturation binding was performed in the same manner, except that the final tracer concentration was 10 nM and the data was collected on a SpectraMax M5 plate reader (Molecular Devices).

#### Competition binding

Peptide competitors were prepared as an 11-point serial dilution at 2X concentration in assay buffer + 2% DMSO. These were then mixed 1:1 with a 2X solution of CHIPOpt tracer (FITC-Ahx-LWWPD) and CHIP protein (43 nM and 20 nM, respectively) to yield a final concentration of 10 nM tracer, 21.5 nM CHIP, and 1% DMSO. An aliquot of the resulting dilutions (18 μL) were then plated and incubated as above, and the resulting polarization data collected on a BioTek H4 plate reader. Raw polarization data (mP) was plotted relative to log_10_[peptide competitor], and data was fit to the log[inhibitor] vs. response model with variable slope in GraphPad PRISM 9.0.

### Peptides & tracers

All peptides were purchased from GenScript as >95% pure N-terminally acetylated peptides unless otherwise indicated. Tracers were prepared identically, with the addition of an N-terminal hexanoic acid linker and FAM fluorophore in place of the acetyl group. Peptide sequences are as follows: IEEVD (Hsp70), MEEVD (Hsp90), IFRPD (CHIC2), LWWPD (CHIPOpt). Synthesis of peptide photo-crosslinkers was performed as previously described^11^, with the following modifications: Following synthesis of the base peptide (M/IEEVD) on Wang resin, an orthogonally protected lysine residue (Fmoc-Lys(Mtt)) was coupled to its N-terminus. MTT deprotection was afforded by incubation with 3% TFA in DCM for 30 minutes at RT. The resin was then washed 3x with DCM, methanol, and DMF. Biotin conjugation was carried out using 1.7 eq. of Biotin-NHS and 20 eq. NMM in 500 uL of DMF while shaking for 1h at RT. Conjugation was then repeated. Following Fmoc deprotection in 4-methylpiperidine, NHS diazirine (Thermo cat. 26167) was attached to the N-terminus using the same conditions. Peptides were cleaved off the resin with 500 μL of cleavage solution (95% trifluoroacetic acid 2.5% water 2.5% triisopropylsilane) while shaking for 1 h, then precipitated in 20 mL cold 1:1 diethyl ether: hexanes. Crude peptides were solubilized in a 1:1:1 mixture DMSO: water: acetonitrile and purified by high-performance liquid chromatography (HPLC) on an Agilent Pursuit 5 C18 column (5 mm bead size, 150 × 21.2mm) using an Agilent PrepStar 218 series preparative HPLC. The mobile phase consisted of A, water 0.1% trifluoroacetic acid and B, acetonitrile 0.1% trifluoroacetic acetic acid. Solvent was removed under reduced atmosphere and purity >95% was confirmed by liquid chromatography-mass spectrometry (LCMS). DMSO stocks were prepared at a concentration of 10 mM based on the gross peptide mass.

### Protein purification

CHIP, CHIP K30A, HOP, and Tau were purified as previously described^11,45,46^. CHIC2 (human, His-tagged) was expressed from a pMCSG7 vector with an N-terminal tobacco etch virus-(TEV)-cleavable 6His Tag. Palmitoylated cysteines (C88, C90, and C92-96) were mutated to serine as previously described^17^ (C6S mutant) to enable expression in Rosetta BL21 (DE3) cells. *E. coli* were grown in terrific broth (TB) at 37 °C, induced with 1mM IPTG in log phase, and grown for 3 hrs at 37 °C. Cells were collected and stored as frozen pellets at −80 °C. Pellets were resuspended in binding buffer (50 mM HEPES pH 7.4, 150 mM NaCl, 5 mM TCEP) supplemented with protease inhibitors (Roche cOmplete), sonicated, clarified, and supernatant was bound to Ni-NTA His-Bind Resin (Novagen) for 1hr at 4 °C. Resin was washed with ∼10 column volumes of binding buffer, then protein was eluted from the resin with His elution buffer (50 mM HEPES pH 7.4, 150 mM NaCl, 5 mM TCEP, 500 mM imidazole). N-terminal His tag was removed by overnight dialysis with TEV protease at 4 °C. Digested material was applied to His-Bind resin to remove cleaved His tag, undigested material and TEV protease. Protein was further purified by size exclusion chromatography (SEC) (Superdex 75) in 50 mM HEPES pH 7.4, 150 mM NaCl, 5mM TCEP and 10% glycerol.

### Protein production for X-ray crystallography

Human CHIP_21-154_-TPR domain was expressed and purified as previously described^47^. HsCHIC2_154-165_-c-term peptide at >95% purity was purchased from Lifetein. Lyophilized HsCHIC2_154-165_ -c-term peptide was dissolved in 25 mM HEPES pH 7.5 and 50 mM NaCl. Briefly, HsCHIP_21-154_-TPR was amplified by PCR from the full-length human CHIP construct and cloned into the pHis||2 expression vector to encode for HsCHIP_21-154_-TPR with an amino-terminal His_6_ tag and an intervening TEV protease cleavage site. The HsCHIP_21-154_-TPR plasmid was transformed into Rosetta2(DE3) *Escherichia coli* competent cells. Expression cultures in Terrific Broth (Fisher BioReagents) were grown at 37 °C until OD_600_ reached 1.0, cooled on ice for 15 minutes, and induced by addition of 400 µM isopropyl β-D-1-thiogalactopyranoside (IPTG). Growth was continued for 20 hours at 18 °C after addition of IPTG prior to harvesting by centrifugation for 10 min at 8,000 × *g*. Frozen cells suspensions were thawed and lysed overnight with slow rotation at 4 °C. Lysate was clarified by centrifugation at 17,000 × *g* for 45 minutes, followed by filtration of lysate supernatant through a 0.45µM filter (Fisher Scientific). His_6_-HsCHIP_21-154_-TPR was purified by Ni2+-affinity chromatography using a 5mL HisTrap HP column (GE Healthcare) loaded in 25 mM HEPES pH 7.5 and 50 mM NaCl. Nonspecifically bound proteins were washed away with 50 mM imidazole and His_6_-HsCHIP_21-154_-TPR eluted in 500 mM imidazole. TEV protease was added at a protease:target protein ratio of 1:20 (w/w), and the mixture was incubated overnight in the presence of 5 mM β-mercaptoethanol. The protein mixture was loaded across a 5mL HisTrap HP column and the flowthrough was collected for further purification by size exclusion chromatography on a HiLoad 16/60 Superdex 75 column. Fractions were analyzed by SDS-PAGE and fractions containing pure HsCHIP_21-154_-TPR were concentrated, frozen drop-wise in liquid nitrogen, and stored at −80 °C.

### CRISPR KO cell line generation

Knockouts were generated using the Synthego Gene Knockout Kit v2 according to the manufacturer’s protocol. Briefly, 500K cells were electroporated with Cas9-RNP complexes using a Lonza nucleofector kit (Program T-016 for K562, T-020 for U251) and grown in 6-well dishes. 72h after nucleofection, pooled knockouts were collected and assayed by western blot. Genomic DNA was extracted using the Qiagen DNEasy kit and PCR of the target loci analyzed using the Synthego ICE tool to confirm high knockout efficiency. Following knockout confirmation, clonal lines were isolated according to Giuliano et al.^48^, then screened by western blot. Selected clones were again genotyped by PCR and ICE analysis (see Extended Data 4), and confirmed mycoplasma negative (ATCC assay) prior to banking.

### Western blotting

Adherent cells were collected by scraping in ice-cold PBS, while centrifugation for 5 min @ 300xg was used for suspension cells. The resulting pellet was washed 1x with PBS, then snap frozen for storage or lysed directly in RIPA buffer (50 mM Tris pH 8.0, 150 mM NaCl, 1% Nonidet P40, 0.5% sodium deoxycholate, 0.1% SDS) supplemented with Roche cOmplete protease inhibitor cocktail. Lysates were incubated for 30 min on ice, then centrifuged at 21,000xg for 10 minutes at 4 °C. The soluble fraction was quantified by BCA assay and normalized to a concentration of 2 mg/mL, then mixed with 3X reducing Laemmli buffer and denatured for 5 minutes at 95 °C. Protein (20 µg) was loaded onto a 4-20% mini-TGX Stain-Free gel, and separated at 200V for 35 mins. For endogenous CHIC2 blots, which required more input for reliable protein detection, 50 to 100 µg material was loaded and the gel was stacked at 60V for 30 mins, followed by separation at 160V for 45 mins. Stain free gels were then activated for 45s using a Chemidoc imager (Bio-Rad), and transferred to 0.2 μm nitrocellulose membranes using the Bio-Rad Trans-Blot Turbo system. Blots were blocked in Intercept TBS blocking buffer (LI-COR) for 30 mins at RT, then incubated with primary antibody in blocking buffer overnight at 4 °C. The following day, blots were washed three times for 5 min each with TBS + 0.05% tween-20 (TBS-T), then incubated with 1:10,000 secondary antibodies (LI-COR) for 1h at RT. Blots were then washed for 3×5 min in TBS-T and imaged on a LI-COR Fc imaging system.

### Immunoprecipitation

293T cells were seeded at a density of 500K cells / well in a 6-well plate (Corning 3335), grown overnight, then transfected using Lipofectamine-3000 according to the manufacturer’s protocol. The following day, cells were harvested in ice-cold PBS, then lysed in ice-cold NP-40 lysis buffer (25 mM Tris-HCl pH 7.4, 150 mM NaCl, 1mM EDTA, 1% NP-40, 5% glycerol, Roche cOmplete protease inhibitor cocktail) by trituration followed by incubation on ice for 10 minutes. Lysates were centrifuged for 10 mins at 21,000xg, and the soluble fraction was harvested and quantified by BCA assay. An input sample was retained, and 100 ug of lysate was then diluted to a final volume of 500 µL in IP lysis buffer. Diluted lysate was added to 20 µL of anti-myc magnetic resin (Pierce cat. 88843) or anti-FLAG magnetic resin (Millipore M8823) in a 1.5 mL low-binding tube (Eppendorf), then incubated at RT for 3 hours with end-over-end rotation. Following incubation, beads were washed with 3×500 µL of IP lysis buffer. Proteins were then eluted by heating to 95 °C in 50 µL of 1X Laemmli buffer for 5 mins, and processed for Western blotting as above.

### *In vitro* ubiquitination

Assays were conducted by preparing a solution of 50 nM E1, 500 nM E2, 125 μM ubiquitin, 500 nM CHIP, 1.25 mM ATP / Mg, and 500 nM substrate in ubiquitination buffer (50 mM Tris, 50 mM KCl, pH 8.0). Reactions were prepared by generating 4 separate 4X stocks of E1/E2/ubiquitin, Ligase/Substrate, CHIC2 / peptide, and ATP+Mg^2+^ in ubiquitination buffer. Protein stocks were combined in equal ratios and equilibrated at RT for 10 minutes, followed by addition of the ATP+Mg^2+^ stock and mixing by pipette to ensure homogeneity. Aliquots of each reaction were removed at the indicated timepoints and quenched in 3X Laemmli buffer. E1 (E-304), UBE2D1 (E2-616), UBE2D1-Ub (E2-800), and Ubiquitin (U-100H) were all purchased from R&D Biosystems.

### Competitive chemoproteomics assays

#### SILAC lysate prep

HEK293T cells were cultured in SILAC DMEM (Thermo) supplemented with 10% dialyzed FBS (Gibco), 483 μM light or heavy Arginine (Sigma A8094 or Cambridge Isotope CNLM-539), and 1 mM light or heavy lysine (Sigma L9037 or Cambridge IsotopeCNLM-291). Cell pellets were harvested and washed 2x with ice-cold PBS, then lysed on ice for 10 minutes in M-PER (Thermo) supplemented with Roche cOmplete protease inhibitor. Lysates were centrifuged at 21,000xg for 10 mins at 4 ℃ and the soluble fraction quantified by BCA assay, then normalized to a concentration of 2 mg/mL. Lysates were then aliquoted, snap frozen, and stored at −80 ℃ for future use.

#### Sample prep & crosslinking

For each sample, 0.5 mg each of light and heavy proteome was pipetted into separate wells of a 96-well PCR plate in 5 × 50 μL aliquots. 2X solutions of crosslinking probe (10 μM each Hsp70 + Hsp90) +/− peptide competitor (100 μM) were prepared in M-PER at a concentration of 2% DMSO. Each peptide solution (5 × 50 µL) was then mixed 1:1 with the aliquoted lysate in the PCR plate (light lysate - competitor, heavy lysate + competitor) to yield a final solution of 1 mg/mL proteome, 5 μM Hsp70/90 probes, +/− 50 μM competitor, 1% DMSO. The resulting samples were incubated for 15 minutes at RT in the dark, then crosslinked under 365nm UV light in the PCR plate for 10 minutes using a 48W nail curing lamp (Sun X9 plus). No-UV controls were prepared as a single 250 μL aliquot in an amber 1.5mL tube (Eppendorf). Following crosslinking, light and heavy aliquots for each sample were pooled in a single 15 mL falcon tube to yield 1 mg of total proteome per sample.

#### Protein enrichment & digestion

Pre-chilled methanol (−30 ℃; 13 mL) was added to each sample, followed by incubation at −30 ℃ overnight to allow complete precipitation to occur. The next day, proteomes were pelleted for 10 mins at 3000xg at 4 ℃. The resulting pellet was washed with 2×1 mL of chilled 1:1 methanol:chloroform, then the pellet was resuspended in 3 mL cold 4:1 methanol:chloroform and centrifuged. The pellet was briefly air-dried, resuspended in 500 μL of freshly prepared 6M Urea, 0.2% SDS in PBS, then tip sonicated for 2×30 s at 40% power to ensure complete re-solubilization. Resuspended proteins were diluted with 1 mL of 0.2% SDS in PBS (2M Urea final), then added to 200 μL of Pierce magnetic streptavidin resin (cat. # 8817) in a 2 mL low-binding tube (Eppendorf). Proteins were captured by continuous rotation at room temperature for 1.5 hours. Using a magnetic rack, beads were then washed with 2×1.75 mL of 0.2% SDS, changing to a new low-binding tube after each wash. Beads were then washed with 2×1.75 mL PBS, then 1.75 mL LC-MS water, and transferred to a new tube during the final wash. Beads were resuspended in 100 μL of 8M Urea in 100 mM TEAB, mixed well, then reduced by addition of 4 μL of 0.5 M TCEP and incubation for 30 min at RT. Cysteines were alkylated with 8 μL of 0.5 M iodoacetamide at RT in the dark for 30 min, and residual iodoacetamide was quenched by addition of 4 μL 1 M DTT. Urea was diluted to 1 M by addition of 700 μL of 100 mM TEAB, then 2 μg of MS-grade trypsin/lys-C mix was added for on-bead digestion overnight at RT with end-over-end mixing. The following day, supernatants were removed and acidified with 200 μL of 10% TFA, then desalted on Thermo SOLA SPE cartridges (cat # 03-150-391). Desalted peptides were dried on a Speedvac, resuspended in 2% acetonitrile with 0.1% formic acid with sonication, and analyzed by LC-MS/MS.

#### LC-MS/MS analysis

1 μg of peptides were injected onto a Thermo Scientific EASY-Spray C18 column (150 mm length, 75 μm diameter, 3 μm particle size) attached to a Dionex UltiMate 3000 NanoRSLC UHPLC. Separation was achieved using a 30-minute linear gradient from 3-40% acetonitrile / 0.1% formic acid at a flow rate of 300 nl/min, followed by a ramp to 80% acetonitrile for column wash prior to re-equilibration. Spectra were acquired on a Thermo Scientific Q-Exactive+ mass spectrometer running a top-12 method. MS1 spectra were acquired from 350-1500 m/z at a resolution of 70,000, with an AGC target of 3e6 and a maximum injection time of 180 ms. MS2 spectra were acquired at a resolution of 35,000 with an isolation width of 1.7 m/z, AGC target of 2e5, maximum injection time of 180 ms, and normalized collision energy of 27. Dynamic exclusion was set to 20s. Data was searched against a non-redundant human proteome database (Uniprot, downloaded 10/2019) using MaxQuant version 1.6.7. N-terminal acetylation and M oxidation were specified as variable modifications, and cysteine carbamidomethylation as a fixed modification. Heavy arginine and lysine modifications were specified as Arg10 and Lys8, respectively. For quantification, min ratio count was set to 2, unique and razor peptides were allowed, and the “re-quantify” option was turned on. All other parameters were set as default. For quantitative analysis of TPR protein binding (as in Fig 2f), data were imported into Skyline for further refinement. A spectral library was generated from the MaxQuant search results, and peptides imported using a FASTA file containing all human CC-TPR proteins. Integration boundaries were manually inspected to ensure consistency across samples, and peptides exhibiting interferences or low quality spectra were excluded. The resulting light/heavy ratios were then exported and plotted in GraphPad PRISM 9.0.

### TMT quantitative proteomics

#### Sample preparation

U251 cells were plated in 6-well plates at a density of 750K cells per well and grown until confluent, then harvested by scraping in ice-cold PBS. K562 cells were plated in 12-well plates at a density of 500K / mL, grown for 24h, then harvested by centrifugation. Cells were washed with 3×1 mL of PBS, then snap frozen prior to further processing using a PreOmics iST-NHS kit. Briefly, pellets were thawed and resuspended in 50 μL of LYSE-NHS solution, then incubated at 95C for 10 minutes with intermittent vortexing every 2 minutes. DNA was sheared by bath sonication until lysates appeared clear, then protein concentration was determined by BCA assay and normalized to 1 mg/mL. Each sample (20 µL) was then transferred to a 0.6 mL low-binding tube (Eppendorf) for digestion and labeling. DIGEST solution (20 µL) was added to each sample, and samples were incubated at 37 °C and 500rpm on a thermomixer for 3 hours. Acetonitrile (15 µL, LCMS-grade) was then added to each sample to achieve a concentration of ∼30% v/v, followed by 100 μg of TMT label (5 μL of 20 μg/μL stock). The labeling reaction was carried out at room temperature for 1h at 500 rpm on the thermomixer, then quenched with 6 μL of 10% hydroxylamine (1% final) and incubation for 15 minutes at 500 rpm. STOP solution (40 µL) was then added, and samples were shaken for an additional 1 minute. Samples were pipetted up and down to ensure homogeneity, then pooled into a single tube. The pooled sample was then split between 2 PreOmics iST cartridges for desalting, and the eluates combined and dried on a Speedvac. Desalted peptides were then resuspended in 600 μL of 0.1% TFA with sonication, and half of the sample (∼90 μg) was fractionated using the Pierce high pH reversed phase fractionation kit (cat # 84868). 8 fractions were collected at 12.5, 15, 17.5, 20, 22.5, 25, 50, and 80% acetonitrile and dried to completion via Speedvac. Peptide fractions were resuspended in 10 μL of 2% ACN / 0.1% FA, sonicated, and analyzed by LC-MS/MS.

#### LC-MS/MS analysis

1 μg of peptides were injected onto a Thermo Scientific EASY-Spray C18 column (150 mm length, 75 μm diameter, 3 μm particle size) attached to a Dionex UltiMate 3000 NanoRSLC UHPLC. Separation was achieved using a 120-minute method composed of a linear gradient from 4-24% acetonitrile over 80 minutes at a flow rate of 200 nl/min, followed by a further ramp to 56% acetonitrile over 25 minutes at 300 nl/min, then a final ramp to 80% acetonitrile for column wash prior to re-equilibration. Spectra were acquired on a Thermo Scientific Q-Exactive+ mass spectrometer running a top-15 method. MS1 spectra were acquired from 375-1400 m/z at a resolution of 70,000, with an AGC target of 3e6 and a maximum injection time of 50 ms. MS2 spectra were acquired at a resolution of 35,000 with an isolation width of 0.7 m/z, AGC target of 1e5, maximum injection time of 100 ms, and normalized collision energy of 32. Dynamic exclusion was set to 30s. Samples were searched against a non-redundant human Uniprot database (downloaded 02/2023) using MaxQuant version 1.6.7. N-acetylation and M oxidation were included as variable modifications, and +113.084 Da was included as a fixed cysteine modification, per PreOmics protocol. MS2 TMT intensities were corrected for isotopic impurities according to the manufacturer’s CoA. All other settings were default. Reverse and contaminant matches were removed using Perseus, then protein intensities were median normalized and analyzed for differential expression using NormalyzerDE^49^. Data was then exported for further analysis in R. Only proteins with 2 or more unique peptides were included unless otherwise indicated.

### Flow cytometry

0.5 - 2 million cells were harvested by light dissociation using TrypLE (Gibco cat. 12-605-010) for 5 minutes at 37C followed by quenching with a 5x volume of complete media. Cell pellets were washed with 2mL of ice-cold FACS buffer (PBS (-Ca, Mg), 1% dialyzed FBS, 5 mM EDTA, 0.02% sodium azide), then resuspended in 100 μL of FACS buffer on ice. 2 μL of conjugated antibody (anti CD119-PE, Miltenyi Biotec cat # 130125874) or isotype control (IgG1-PE, Miltenyi Biotec cat # 130113450) was spiked into each sample, briefly vortexed, and incubated in the dark for 10 minutes. The antibody solution was diluted with 1.9 mL FACS buffer, then cells were centrifuged for 5 minutes at 300xg. The resulting pellet was resuspended in 500 μL of FACS buffer, strained, and analyzed directly on a BD LSRFortessa X-14 flow cytometer. FlowJo software was used to select a viable, single-cell population based on forward and side scatter profiles, and IFNGR-positive cells were gated based on the isotype control.

### Cell culture

All cell lines were maintained at 37C and 5% CO_2_. HEK293T, HEK293 Flp-In T-REX, and U251 cells were maintained in DMEM (Gibco cat. # 11995065). K562 cells were maintained in IMDM (Gibco cat. # 12440053). All media was supplemented with 10% HI FBS (Gibco cat # 10438026).

### Immunofluorescence

HEK-293 cells were seeded on poly-lysine coated coverslips in 12-well dishes and grown overnight. The following day, cells were transfected using LTX-3000 according to the manufacturer’s protocol, then grown overnight. The next day, cells were stained according to Stadler et al^50^. Briefly, media was aspirated and cells were fixed using 4% PFA in media (DMEM + 10% FBS) on ice for 15 minutes. Fixed wells were washed 2x with PBS at RT, then incubated overnight at 4C in 500 μL of primary antibody (1 μg/mL Ms anti myc, Invitrogen cat. #13-500) diluted in blocking / permeabilization buffer (PBS + 4% FBS, 0.1% saponin). The next day, primary antibody was aspirated and cells washed in PBS for 4×10 mins at RT. Secondary antibody (Invitrogen anti-MS AF-488, cat # A-11001) was diluted to 1 μg/mL in blocking / permeabilization buffer, and 500 μL added to each well for 1.5 hours at RT. Nuclei were stained with 1 μg/mL Hoescht 33342 in PBS for 5 minutes, then cells were washed with PBS for 4×10 mins at RT. Coverslips were mounted onto slides using ProLong Gold Antifade Mountant (Thermo cat #P10144) and allowed to cure overnight at RT while protected from light. Images were collected on a Leica SP8 confocal microscope, and processed in Fiji.

### Live-cell imaging

HEK-293 Flp-In T-REX cells expressing mEGFP-CHIC2 constructs were generated according to the manufacturer’s instructions (https://www.thermofisher.com/order/catalog/product/R78007). Cells were seeded in 6-well plates, then transfected using LTX-3000 as above. The following day, cells were trypsinized and seeded at a density of 25,000 cells / well in 96-well imaging plates (Greiner cat. 655090), then allowed to grow overnight in media containing 10 ng/mL doxycycline. Prior to imaging the next day, cells were treated for 4 hours with the appropriate compound in complete media (100 nM Bafilomycin A1, 50 nM bortezomib), with the exception of Bromo-palmitate which was added at a concentration of 25 μM immediately upon dox induction the prior day. Following compound treatment, nuclei were stained with 5 μg/mL Hoescht 33342 in HBSS for 5 minutes. Cells were washed with HBSS, then returned to complete media for imaging. Images were collected on an IN Cell 2000 confocal microscope and analyzed in Fiji.

### Crystallization, X-ray crystallographic diffraction data collection, and structural refinement

Crystals of the CHIP_21-154_-TPR:CHIC2_154-165_-c-term complex were obtained by sitting drop vapor diffusion in 96-well IntelliPlates (Art Robbins) set up with a Phoenix crystallization robot (Art Robbins). Protein and peptide were combined at 7 mg/mL concentration with a 3:1 peptide:protein ratio) was mixed with crystallization conditions from MCSG1-4 (Microlytic) and BCS sparse matrix crystallization screens at volumes of 400 nL CHIP/CHIC2 and 400 nL condition. Crystals of were harvested and frozen in liquid nitrogen within one month of setting up sitting drop vapor diffusion trials. Crystals were frozen in liquid nitrogen 60-90 days after setting up sitting drop vapor diffusion trials. The CHIP_21-154_-TPR:CHIC2_154-165_-c-term complex was crystallized in 0.05 M Magnesium sulfate heptahydrate 0.1 M HEPES pH 7.5, and 28 % v/v PEG Smear Medium. Harvested crystals were swished through LV CryoOil (MiTeGen) for cryoprotection immediately prior to freezing in liquid nitrogen. X-ray diffraction data were collected on beamline 4.2.2 at 1.000020 Å wavelength at the Advanced Light Source at Lawrence Berkeley National Laboratory. X-ray diffraction data were processed in XDS^51^, followed by molecular replacement with PHASER^52^ using our prior CHIP-TPR structure (PDB ID 4KBQ, chain A) as the molecular replacement search model. Model building and refinement were performed iteratively using PHENIX^53^ and Coot^54^. Coordinates and experimental data for the CHIP_21-154_-TPR:CHIC2_154-165_-c-term complex were deposited in the PDB with accession code 8SUV. Geometrical and stereochemical validation was conducted using MolProbity^55^. All figures were prepared using UCSF Chimera^56^.

### C. elegans Assays

#### Strains

The following strains were used for experiments described in this manuscript: N2 (Bristol) wild-type, *chn-1* (*by155*) I, *tag-266* (*ok2462*) III, *tag-266* (*syb5138*) III, *wdIs51* (*P_F49H12.4_::GFP*). All strains were maintained at 20 °C as described previously^57^.

#### Longevity

Briefly, a timed egg lay was performed to gather an age-synchronized population of animals. Animals were plated on 60-mm NGM agar plates containing a thin lawn of OP50 *Escherichia coli* spotted the day before the experiment. Each day, or every other day, the number of living and dead worms were recorded until there were no living animals remaining. Longevity assays were performed using 75 animals over 3 independent experiments by an experimenter blinded to the genotypes of the animals being tested.

#### Quantification of PVD Degeneration

Fluorescence imaging of *wdIs51* (*P_F49H12.4_::GFP*) was performed as follows. *wdIs51* expresses GFP in the neuronal pair PVDR and PVDL, which have extensive dendritic branching. Quantification of PVD neurodegeneration was performed as described^31^. Briefly, Larval-stage 4 (L4) animals were immobilized in a droplet of M9 containing 2.5 mM Levamisole (Tetramisole, Sigma) and placed on a 2% agarose pad. PVD neurons were imaged using 20× magnification on a Zeiss Axio Imager M2. Animals were scored for PVD neuron branching and bead-like puncta in dendrites. Degeneration assays were performed using 30 animals over 3 independent experiments by an experimenter blinded to the genotypes of the animals being tested.

### Antibodies

Rb CHIP (Abcam EPR4447, 1:2000), Ms CHIC2 (sc-515175, 1:500), Rb Myc (sc-789, 1:500), Ms Hsp70 (SCBT sc-137239, 1:500), Ms Hsp90 (SCBT sc-13119, 1:500), Rb STIP1/HOP (Abcam EPR6605, 1:2000), Ms Tau5 (Invitrogen AHB0042,, 1:1000), Ms Ubiquitin (CST 3936, 1:2000), Rb UBE2D1 (Invitrogen PA576645, 1:2000), Rb FLAG (CST 14793, 1:2000), Rb GAPDH (CST 2118S, 1:2000), Ms Tubulin (CST 2144S, 1:2000), Ms IFNGR (sc-28363, 1:500), anti CD119-PE (Miltenyi Biotec 130125874), IgG1-PE (Miltenyi Biotec 130113450).

## Data availability

<u>Raw proteomic data have been deposited to the ProteomeXchange consortium via the PRIDE partner repository with the identifiers</u> PXD043803 (TMT proteomics of CHIC2 and CHIP knockout) and PXD043804 (competitive chemoproteomics of TPR cochaperone proteins). Processed proteomic data are provided in the supplementary information.

## Supplemental methods

**Supplemental Table S1:** Data collection and refinement statistics for VIM-20 crystal structures.

|  |  |
| --- | --- |
|  | CHIP-TPR:CHIC2 |
| Data Collection |  |
| X-ray source | Beamline ALS 4.2.2 |
| Wavelength (Å) | 1.000020 |
| Resolution range (Å)<sup>b</sup> | 46.53 – 1.63 (1.688 – 1.63) |
| Space group | *P* 1 2<sub>1</sub> 1 |
| *a*, *b*, *c* (Å) | 46.53, 82.36, 78.39 |
| *a*, *b*, *g* (°) | 90, 90.03, 90 |
| Reflections<sup>b</sup> | 498,239 (50,246) |
| Unique reflections<sup>b</sup> | 71,293 (7,022) |
| Multiplicity<sup>b</sup> | 7.0 (7.2) |
| Completeness (%)<sup>b</sup> | 96.4 (94.6) |
| Mean *I*/sigma (*I*)<sup>b</sup> | 8.52 (0.88) |
| Wilson B-factor (Å<sup>2</sup>) | 20.6 |
| CC<sub>1/2</sub><sup>b</sup> | 0.998 (0.652) |
| Refinement |  |
| PDB ID | 8SUV |
| Number of reflections<sup>b</sup> | 71,065 (6,976) |
| Reflections used for R<sub>free</sub><sup>b</sup> | 1,540 (136) |
| *R*<sub>work</sub> / *R*<sub>free</sub> | 0.274 / 0.290 |
| Number of atoms<br>(protein/ligands/water) | 4,395 / 20 / 404 |
| Average *B* factors<br>(protein/ligands/water) | 32.5 / 37.7 / 31.2 |
| Model Quality |  |
| Bond length rmsd (Å) | 0.012 |
| Bond angle rmsd (°) | 1.550 |
| Ramachandran favored (%) | 98.1 |
| Ramachandran allowed (%) | 100 |
| Ramachandran outliers (%) | 0 |
| Poor rotamers (%)<sup>b</sup> | 0.22 |
| Clashscore<sup>b</sup> | 4.79 |
| Clash percentile<sup>b</sup> | 94<sup>th</sup> percentile<br>(N=1,784; all resolutions) |
| MolProbity score<sup>b</sup> | 1.25 |
| MolProbity score percentile<sup>b</sup> | 99<sup>th</sup> percentile<br>(N=27,675; all resolutions) |

<sup>a</sup> Data for the highest resolution shell is represented by the values in parentheses within the table.
<sup>b</sup> Values calculated using MolProbity v4.5.2<sup>55</sup>.

## Notes

### Competing Interest Statement

The authors have declared no competing interest.

## References

1. Rape, M. Ubiquitylation at the crossroads of development and disease. Nat. Rev. Mol. Cell Biol. 19, 59–70 (2018).

2. Lorenz, S., Cantor, A. J., Rape, M. & Kuriyan, J. Macromolecular juggling by ubiquitylation enzymes. BMC Biol. 11, 65 (2013).

3. Haakonsen, D. L. & Rape, M. Branching Out: Improved Signaling by Heterotypic Ubiquitin Chains. Trends Cell Biol. 29, 704–716 (2019).

4. Herhaus, L. & Dikic, I. Expanding the ubiquitin code through post-translational modification. EMBO Rep. 16, 1071–1083 (2015).

5. Balaji, V. & Hoppe, T. Regulation of E3 ubiquitin ligases by homotypic and heterotypic assembly. F1000Research 9, 88 (2020).

6. Miller, J. J. et al. Emi1 stably binds and inhibits the anaphase-promoting complex/cyclosome as a pseudosubstrate inhibitor. Genes Dev. 20, 2410–2420 (2006).

7. Qian, S.-B., McDonough, H., Boellmann, F., Cyr, D. M. & Patterson, C. CHIP-mediated stress recovery by sequential ubiquitination of substrates and Hsp70. Nature 440, 551–555 (2006).

8. Dickey, C. A., Patterson, C., Dickson, D. & Petrucelli, L. Brain CHIP: removing the culprits in neurodegenerative disease. Trends Mol. Med. 13, 32–38 (2007).

9. Ballinger, C. A. et al. Identification of CHIP, a Novel Tetratricopeptide Repeat-Containing Protein That Interacts with Heat Shock Proteins and Negatively Regulates Chaperone Functions. Mol. Cell. Biol. 19, 4535–4545 (1999).

10. Zhang, M. et al. Chaperoned Ubiquitylation—Crystal Structures of the CHIP U Box E3 Ubiquitin Ligase and a CHIP-Ubc13-Uev1a Complex. Mol. Cell 20, 525–538 (2005).

11. Ravalin, M. et al. Specificity for latent C termini links the E3 ubiquitin ligase CHIP to caspases. Nat. Chem. Biol. 15, 786 (2019).

12. Reinhardt, L. et al. Dual truncation of tau by caspase-2 accelerates its CHIP-mediated degradation. Neurobiol. Dis. 182, 106126 (2023).

13. Narayan, V., Pion, E., Landré, V., Müller, P. & Ball, K. L. Docking-dependent Ubiquitination of the Interferon Regulatory Factor-1 Tumor Suppressor Protein by the Ubiquitin Ligase CHIP *. J. Biol. Chem. 286, 607–619 (2011).

14. Nadel, C. M. et al. The E3 Ubiquitin Ligase, CHIP/STUB1, Inhibits Aggregation of Phosphorylated Proteoforms of Microtubule-associated Protein Tau (MAPT). J. Mol. Biol. 435, 168026 (2023).

15. Tsherniak, A. et al. Defining a Cancer Dependency Map. Cell 170, 564–576.e16 (2017).

16. Wang, L. et al. Molecular Mechanism of the Negative Regulation of Smad1/5 Protein by Carboxyl Terminus of Hsc70-interacting Protein (CHIP) *. J. Biol. Chem. 286, 15883–15894 (2011).

17. Cools, J., Mentens, N. & Marynen, P. A new family of small, palmitoylated, membrane-associated proteins, characterized by the presence of a cysteine-rich hydrophobic motif. FEBS Lett. 492, 204–209 (2001).

18. Ng, S. et al. STUB1 is an intracellular checkpoint for interferon gamma sensing. Sci. Rep. 12, 14087 (2022).

19. Haslbeck, V. et al. Chaperone-Interacting TPR Proteins in Caenorhabditis elegans. J. Mol. Biol. 425, 2922–2939 (2013).

20. Taipale, M. et al. A Quantitative Chaperone Interaction Network Reveals the Architecture of Cellular Protein Homeostasis Pathways. Cell 158, 434–448 (2014).

21. Smith, M. C. et al. The E3 Ubiquitin Ligase CHIP and the Molecular Chaperone Hsc70 Form a Dynamic, Tethered Complex. Biochemistry 52, 5354–5364 (2013).

22. Rebeyev, N. CHIC2 and STUB1 regulate interferon-γ receptor cell surface expression. Cambridge University Doctoral Thesis (2019).

23. Apriamashvili, G. et al. Ubiquitin ligase STUB1 destabilizes IFNγ-receptor complex to suppress tumor IFNγ signaling. Nat. Commun. 13, 1923 (2022).

24. Koochaki, S. H. J. et al. A STUB1 ubiquitin ligase/CHIC2 protein complex negatively regulates the IL-3, IL-5, and GM-CSF cytokine receptor common β chain (CSF2RB) protein stability. J. Biol. Chem. 102484 (2022) doi:10.1016/j.jbc.2022.102484.

25. Okiyoneda, T. et al. Peripheral Protein Quality Control Removes Unfolded CFTR from the Plasma Membrane. Science 329, 805–810 (2010).

26. Tawo, R. et al. The Ubiquitin Ligase CHIP Integrates Proteostasis and Aging by Regulation of Insulin Receptor Turnover. Cell 169, 470–482.e13 (2017).

27. Slotman, J. A. et al. Ubc13 and COOH Terminus of Hsp70-interacting Protein (CHIP) Are Required for Growth Hormone Receptor Endocytosis. J. Biol. Chem. 287, 15533–15543 (2012).

28. Min, J.-N. et al. CHIP Deficiency Decreases Longevity, with Accelerated Aging Phenotypes Accompanied by Altered Protein Quality Control. Mol. Cell. Biol. 28, 4018–4025 (2008).

29. tag-266 (gene) - WormBase : Nematode Information Resource. https://wormbase.org/species/c_elegans/gene/WBGene00044319#0-9fb867d124ha-5;class=Gene.

30. chn-1 (gene) - WormBase : Nematode Information Resource. https://wormbase.org/species/c_elegans/gene/WBGene00000500#0-9fb867d124ha-10;class=Gene.

31. Umano, A. et al. The molecular basis of spinocerebellar ataxia type 48 caused by a de novo mutation in the ubiquitin ligase CHIP. J. Biol. Chem. 298, 101899 (2022).

32. Zhang, S., Hu, Z., Mao, C., Shi, C. & Xu, Y. CHIP as a therapeutic target for neurological diseases. Cell Death Dis. 11, 1–12 (2020).

33. Mol, M. O. et al. Clinical and pathologic phenotype of a large family with heterozygous *STUB1* mutation. Neurol. Genet. 6, e417 (2020).

34. Shi, C. et al. Disrupted structure and aberrant function of CHIP mediates the loss of motor and cognitive function in preclinical models of SCAR16. PLOS Genet. 14, e1007664 (2018).

35. Dickey, C. A. et al. Deletion of the Ubiquitin Ligase CHIP Leads to the Accumulation, But Not the Aggregation, of Both Endogenous Phospho- and Caspase-3-Cleaved Tau Species. J. Neurosci. 26, 6985–6996 (2006).

36. Stankiewicz, M., Nikolay, R., Rybin, V. & Mayer, M. P. CHIP participates in protein triage decisions by preferentially ubiquitinating Hsp70-bound substrates. FEBS J. 277, 3353–3367 (2010).

37. Welch, W. J. & Feramisco, J. R. Purification of the major mammalian heat shock proteins. J. Biol. Chem. 257, 14949–14959 (1982).

38. Muller, P. et al. C-terminal phosphorylation of Hsp70 and Hsp90 regulates alternate binding to co-chaperones CHIP and HOP to determine cellular protein folding/degradation balances. Oncogene 32, 3101–3110 (2013).

39. Thul, P. J. et al. A subcellular map of the human proteome. Science 356, eaal3321 (2017).

40. Haglund, K. & Dikic, I. The role of ubiquitylation in receptor endocytosis and endosomal sorting. J. Cell Sci. 125, 265–275 (2012).

41. Balaji, V. et al. A dimer-monomer switch controls CHIP-dependent substrate ubiquitylation and processing. Mol. Cell 82, 3239–3254.e11 (2022).

42. Cools, J. et al. Fusion of a Novel Gene, BTL, to ETV6 in Acute Myeloid Leukemias With a t(4;12)(q11-q12;p13). Blood 94, 1820–1824 (1999).

43. Pardanani, A. et al. CHIC2 deletion, a surrogate for FIP1L1-PDGFRA fusion, occurs in systemic mastocytosis associated with eosinophilia and predicts response to imatinib mesylate therapy. Blood 102, 3093–3096 (2003).

44. Griss, J. et al. ReactomeGSA - Efficient Multi-Omics Comparative Pathway Analysis. Mol. Cell. Proteomics 19, 2115–2125 (2020).

45. Assimon, V. A., Southworth, D. R. & Gestwicki, J. E. Specific Binding of Tetratricopeptide Repeat Proteins to Heat Shock Protein 70 (Hsp70) and Heat Shock Protein 90 (Hsp90) Is Regulated by Affinity and Phosphorylation. Biochemistry 54, 7120–7131 (2015).

46. Montgomery, K. M. et al. Chemical Features of Polyanions Modulate Tau Aggregation and Conformational States. J. Am. Chem. Soc. 145, 3926–3936 (2023).

47. Zhang, H. et al. A Bipartite Interaction between Hsp70 and CHIP Regulates Ubiquitination of Chaperoned Client Proteins. Structure 23, 472–482 (2015).

48. Giuliano, C. J., Lin, A., Girish, V. & Sheltzer, J. M. Generating Single Cell–Derived Knockout Clones in Mammalian Cells with CRISPR/Cas9. Curr. Protoc. Mol. Biol. 128, (2019).

49. Willforss, J., Chawade, A. & Levander, F. NormalyzerDE: Online Tool for Improved Normalization of Omics Expression Data and High-Sensitivity Differential Expression Analysis. J. Proteome Res. 18, 732–740 (2019).

50. Stadler, C., Skogs, M., Brismar, H., Uhlén, M. & Lundberg, E. A single fixation protocol for proteome-wide immunofluorescence localization studies. J. Proteomics 73, 1067–1078 (2010).

51. Kabsch, W. Integration, scaling, space-group assignment and post-refinement. Acta Crystallogr. D Biol. Crystallogr. 66, 133–144 (2010).

52. McCoy, A. J. et al. Phaser crystallographic software. J. Appl. Crystallogr. 40, 658–674 (2007).

53. Adams, P. D. et al. PHENIX: a comprehensive Python-based system for macromolecular structure solution. Acta Crystallogr. D Biol. Crystallogr. 66, 213–221 (2010).

54. Emsley, P., Lohkamp, B., Scott, W. G. & Cowtan, K. Features and development of Coot. Acta Crystallogr. D Biol. Crystallogr. 66, 486–501 (2010).

55. Chen, V. B. et al. MolProbity: all-atom structure validation for macromolecular crystallography. Acta Crystallogr. D Biol. Crystallogr. 66, 12–21 (2010).

56. Pettersen, E. F. et al. UCSF Chimera—A visualization system for exploratory research and analysis. J. Comput. Chem. 25, 1605–1612 (2004).

57. Brenner, S. THE GENETICS OF CAENORHABDITIS ELEGANS. Genetics 77, 71–94 (1974).

